# Genome-resolved expansion of *Nucleocytoviricota* and *Mirusviricota* reveals new diversity, functional potential, and biotechnological applications

**DOI:** 10.1101/2025.09.26.678796

**Authors:** Yumary M. Vasquez, Tiago Nardi, Gbocho Masato Terasaki, Petra Byl, Tomáš Brůna, Juan C. Villada, Miguel F. Romero, Thomas Mock, Timothy Y. James, GVMAGs data consortium, Tanja Woyke, Frederik Schulz

**Affiliations:** United States Department of Energy Joint Genome Institute, Berkeley, CA, USA; Department of Biology and Biotechnology, University of Pavia, Italy; Department of Applied Math, University of California, Merced, Merced, CA, USA; School of Ocean and Earth Science and Technology, University of Hawai‘i Mānoa, Honolulu, HI USA; Instituto de Biología, Universidad Nacional Autónoma de México, 3er. Circuito Exterior S/N, Ciudad Universitaria, CP 04510 CDMX, México; School of Environmental Sciences, University of East Anglia, Norwich Research Park, Norwich, UK; Department of Ecology and Evolutionary Biology, University of Michigan, Ann Arbor, MI USA; Departments of Civil and Environmental Engineering and Bacteriology, University of Wisconsin - Madison, Madison, WI, USA; School of Biotechnology and Biomolecular Sciences, UNSW Sydney, Sydney, Australia; Department of Microbiology & Immunology, University of British Columbia, Vancouver, Canada; Department of Biological Sciences College of Science, Clemson University, SC, USA; College of Earth, Ocean, and Atmospheric Sciences Oregon State University, Corvallis, OR; Microbial Ecology, Duke University, NC, USA; Department of Bacteriology, University of Wisconsin-Madison, Madison, WI, USA; Division of Environmental and Biomolecular Systems, Institute of Environmental Health, Oregon Health & Science University, Portland, OR, USA; Great Lakes Institute for Environmental Research, University of Windsor, Ontario Canada; Biology Department, Concordia, University, Montreal Canada; Department of Biology, Kenyon College, Gambier, OH, USA; School of Civil & Environmental Engineering, Georgia Tech, Atlanta, GA, USA; Department of Earth Science and Marine Science Institute, UC Santa Barbara, Santa Barbara, CA, USA; Environmental Genomics and Systems Biology, Lawrence Berkeley National Lab, Berkeley CA, USA; Department of Biology and Department of Estuarine and Ocean Sciences University of Massachusetts Dartmouth, MA, USA; Department of Chemistry and Biochemistry, Thermal Biology Institute, Montana State University, Bozeman, MT, USA; College of Arts and Sciences and College of Engineering Oregon State University Columbus, OH, USA; Departments of Microbiology and Immunology and Earth, Ocean, and Atmospheric Sciences University of British Columbia, Vancouver, Canada; Clark University, Lasry Center for Bioscience, Worcester, MA, USA

## Abstract

Nucleocytoplasmic large DNA viruses (NCLDV) of the phyla *Nucleocytoviricota* and viruses of the newly proposed *Duplodnaviria* phylum, *Mirusviricota*, exhibit taxonomic richness which continues to expand due to metagenomic sequencing of Earth’s biomes. Giant viruses contain complex genomes encoding genes of both viral and cellular origin, representing a reservoir of unexplored biological functions with potential implications for ecology, evolution, and biotechnology. Here, we present the largest curated database of giant virus metagenome-assembled genomes (GVMAGs V2), comprising 8,508 species-level clusters inferred from 18,727 genomes, originating from marine, freshwater, anthropogenic and terrestrial environments, a six-fold increase from the previous giant virus phylogenetic frameworks. Phylogenomics and relative evolutionary distance analysis revealed 712 novel genera, 13 previously unknown viral families and a new proposed order, tentatively named *Mycodnavirales.* We improved gene calling of 12% of giant virus genomes by accounting for alternative and custom genetic codes, enabling more accurate identification of protein-coding genes. Database mining uncovered endogenous viral elements in a broad spectrum of eukaryotes, spanning algae, fungi, and parasitic protists highlighting that giant virus integration is both widespread and evolutionarily persistent. Orthologous clustering of 2.5 million proteins identified 135,998 orthogroups representing comprehensive metabolic capabilities, such as enrichment of genes involved in aromatic compound degradation (commonly associated with bioremediation) in *Algavirales* genomes. Furthermore, we detected widespread biosynthetic gene clusters underpinning antimicrobial activity and antibiotic resistance, suggesting roles of giant viruses in host defense and in the dissemination of antibiotic resistance genes. Conversely, 67% of orthogroups have unknown functions, underscoring a substantial unexplored potential. This comprehensive publicly available database provides a critical resource for the giant virus research community and a foundation for uncovering virus-host interactions, exploring viral evolution, and identifying reservoirs for novel enzymes with the potential to advance biotechnological applications.

## MAIN

Nucleocytoplasmic large DNA viruses (*Nucleocytoviricota*), commonly referred to as giant viruses (GVs), represent a diverse group of viruses that challenge traditional virological paradigms due to their large size, genomic complexity, and unique biological features (Raoult *et al*., 2004; Abergel *et al*., 2007; Yutin *et al*., 2013; Schulz *et al*., 2017). Their particles rival small bacteria in size and their genomes can exceed 2.5 Mb, encoding hundreds to thousands of proteins, some drawn from cellular translation and metabolic pathways (Raoult *et al*., 2004; Philippe *et al*., 2013; Schulz, Abergel and Woyke, 2022). Metagenomic surveys have discovered giant viruses across Earth’s major biomes, suggesting they have a significant yet still poorly understood ecological role (Schulz, Abergel and Woyke, 2022).

Despite these metagenomic surveys, the taxonomic and functional landscape of *Nucleocytoviricota* remains fragmentary because of sampling biases towards marine and freshwater ecosystems and isolated algae and protists (Legendre *et al*., 2018; Mihara *et al*., 2018; Moniruzzaman, Weinheimer, *et al*., 2020; Schulz *et al*., 2020; Aylward *et al*., 2021, 2023).

Existing giant virus genome catalogues have laid groundwork for *Nucleocytoviricota* research, yet they still encompass only a fraction of the group’s diversity and were generated with gene-calling pipelines that assume the standard genetic code for all genomes. Consequently, some open reading frames remain unannotated and occasional cellular sequences may fail quality filters. Expanding *Nucleocytoviricota* metagenome resources with new sampling ecosystems and with better protein prediction will lead to a more comprehensive comparative analysis. Additionally, these resources can be used to clarify lineage-specific metabolic innovations and increase our understanding of *Nucleocytoviricota* diversity and evolution.

Here we have assembled GVMAGs V2, the largest curated collection of giant virus metagenome-assembled genomes (GVMAGs) to date, a four-fold increase from the original GVMAGs database (Schulz et al., 2020) and a six-fold increase from the previous 1,382 species-level representatives (Aylward et al., 2021). We screened public assemblies in IMG/M and integrated data from additional sources, recovering 18,727 giant virus genomes (Benson *et al*., 2012; Tully *et al*., 2017; Bäckström *et al*., 2019; Moniruzzaman, Martinez-Gutierrez, *et al*., 2020; Aylward *et al*., 2021; Chen *et al*., 2023; Gaïa *et al*., 2023; Oliver *et al*., 2024; Schulz *et al*., 2024; Rohwer *et al*., 2025). De-replication at 95% average nucleotide identity resulted in 8,508 species-level clusters, and resolved 712 previously unrecognised genera. Subsequent phylogenomic analysis supported the proposal of 13 new viral families, markedly expanding the current taxonomy. Additionally, we propose a new order (*Mycodnavirales*) which is basal to *Imitervirales,* and incorporate genomes from the recently described *Duplodnaviria* phylum, *Mirusviricota* (Gaïa *et al*., 2023), into our comparative analyses, extending the evolutionary and functional scope of the dataset.

To enhance genome annotation accuracy, we accounted for alternative genetic codes during gene prediction. We increased more complete profile hits and higher average scores in 1,066 genomes, indicating an improvement in annotation quality and recovering >1300 previously missed genes. We mined PhycoCosm, MycoCosm and IMG/M databases to recover endogenous viral elements in eukaryotic genomes (Grigoriev et al., 2014, 2021; Chen et al., 2019). Ortholog clustering of 2.5 million proteins resolved 135,998 gene families, only 33% of which have functional assignments, revealing a significant reservoir of unknown biology. Module-level pathway reconstruction revealed order- and family-specific enrichment for photosynthesis, formaldehyde assimilation and aromatic degradation offering new insights into virus-driven metabolic reprogramming. In addition, we identified widespread biosynthetic gene clusters and antibiotic resistance genes within giant virus genomes, highlighting their potential as reservoirs and vehicles for novel enzymes, bioactive compounds, and resistance determinants of biotechnological interest. To enable easy exploration of these data, we provide an interactive, agentic system (https://newlineages.com/pages/tools/gv-ai-agent). Thus, GVMAGs V2 provides a phylogenetic and functional resource for enabling comparative analyses of giant virus evolution, ecology, and biotechnological potential.

## RESULTS & DISCUSSION

### New families, genera, and order from an expanded giant virus genome database

Expanding the genomic catalog of giant viruses is essential for improving taxonomic resolution and uncovering novel lineages. We collated a total of 18,727 genomes from multiple data sources, including genomes recovered from the IMG/M database (Chen *et al*., 2023; Villada *et al*., 2025), a co-assembly and single assembly of MAGs from the freshwater Lake Mendota (Rohwer *et al*., 2023; Oliver *et al*., 2024), genomes from the giant virus taxonomy framework (Moniruzzaman, Martinez-Gutierrez, et al., 2020; Aylward et al., 2021), TARA Oceans sequencing efforts (Tully *et al*., 2017; Gaïa *et al*., 2023), isolate giant virus genomes (Benson *et al*., 2012), a recent protists single cell genome study (Schulz et al., 2024), and from deep sea sediments (Bäckström et al., 2019) (Figure 1a). First, dereplication using Average Nucleotide Identity (ANI) resulted in 8,508 non-redundant species-level genomes (Roux *et al*., 2019), separated into 2,484 clusters and 6,024 singletons (hereafter known as species-level-clusters; see Methods for more information) (Figure 1b, Supplemental Table 1). The species-level-clusters indicated here represent a four-fold increase from the original GVMAGs database (Schulz et al., 2020) and a six-fold increase from the previous 1,382 species-level representative framework (Aylward et al., 2021). Following phylogenetic tree building, 469 genomes were removed, including genomes that contained markers shorter than 50% of the marker median length. Using a phylogenetic based-pairwise distance matrix (PDM), genomes were further clustered into 851 clusters and 1,756 singletons (hereafter known as PDM-clusters; see Methods for more information) (Figure 1c). Lastly, filtering for a minimum number of giant virus orthologous groups (GVOGs), the final phylogenetic tree contained 2,037 genomes, including the addition of 4 *Proculovirales* genomes and 192 *Egovirales* genomes (Gaïa *et al*., 2024) (Figure 1d). This phylogenetic tree was created using a concatenated alignment of seven GVOG markers identified in Aylward et al. (2021) (hereafter identified as GVOG7 PDM-clustered tree). This tree does not include *Mirusviricota* genomes as they are missing most of the *Nucleocytoviricota* hallmark genes.

**Figure 1:**
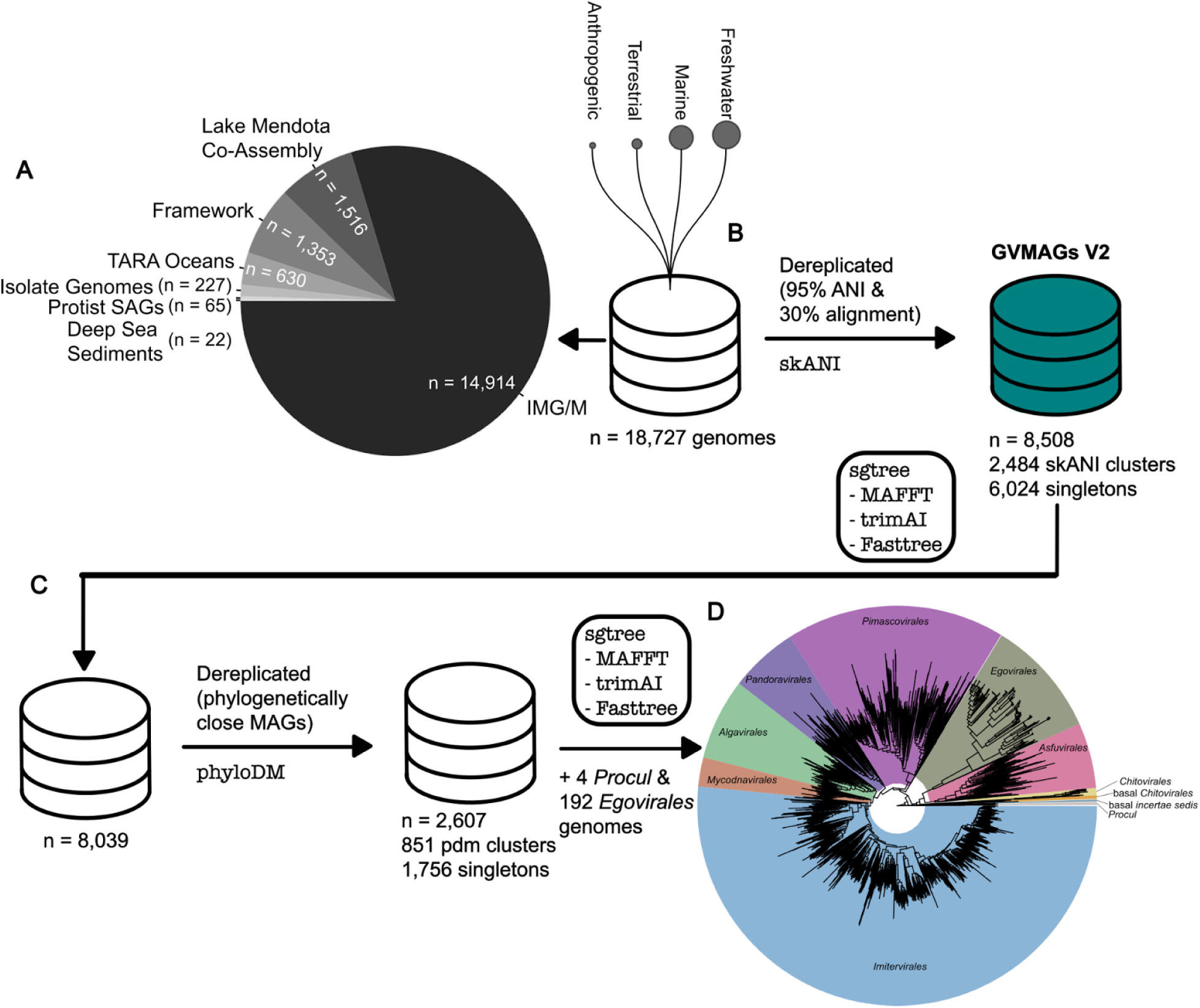
Workflow for genomic dereplication. A total of 18,727 genomes were obtained from public resources, predominantly from the IMG/M database (A). The workflow consisted of three main stages: B) Initial dereplication using skANI (sequence-based average nucleotide identity) with thresholds of 95% ANI and 30% minimum alignment coverage, generating preliminary clusters each represented by a single genome representing the GVMAGs V2 database; C) Construction of an initial phylogenetic tree, followed by secondary dereplication based on phylogenetic distance matrix (PDM) analysis of closely related MAGs (metagenome-assembled genomes), resulting in 2,607 representative genomes and; D) Incorporation of an additional 196 genomes in the final phylogenetic tree.

Using the GVOG7 PDM-clustered tree, we quantified relative evolutionary distance (RED) scores ranging from 0 to 1, where groups with scores closer to zero represent deeper evolutionary lineages near the root, while those with scores closer to one indicate narrowly defined clades positioned near the terminal branches. The RED scores of the *Nucleocytoviricota* classes range from 0.18 to 0.2, whereas the values for the orders range from 0.26 to 0.32, families range from 0.35 to 0.75 and genera range from 0.76 to 0.98 (Supplemental Figure S1, Supplemental Table 2). To prevent misclassifications based on isolated genome placements, we assigned family taxonomy if at least three genomes were present in the taxa, otherwise unclassified families were labeled with “incertae sedis”. Using this method, we recovered a total of 61 families, including 33 “incertae sedis” cluster representatives, which likely represent additional families with currently insufficient sampling, and 1,030 genera. Using our expanded giant virus database, we identified new families and genera using RED score analysis on our species tree that was inferred from representative genomes obtained after PDM clustering of viral species. For newly proposed families and genera, we assigned nonredundant identifiers following previous nomenclature (Aylward *et al*., 2021). We identified 13 new family classifications within the entire *Nucleocytoviricota,* none of which contain single cultivated representatives. *Imitervirales*, the largest giant virus order, displayed five new family classifications, a notable increase from the 11 families identified in 2021 (Aylward *et al*., 2021) (Figure 2). New family classifications were also identified in *Pimascovirales* (n = 4), *Pandoravirales* (n = 3) and one new family within *Asfuvirales*. Most newly proposed families are a mixture of clustered and singleton genomes, with the exception of genomes in the newly proposed families: IM_24 and MY_01. These families only contain singletons with no species-level- or PDM-clusters, possibly indicating new genera and species that have yet to be fully described and require further sampling to confirm. At the genus level, we identified 712 new genera. Of these, 343 are singletons represented by only one genome. The remaining 369 genera contained 1,092 species-level clusters and 3,767 genomes (taking into account all strain genomes in the species-level clusters).

**Figure 2:**
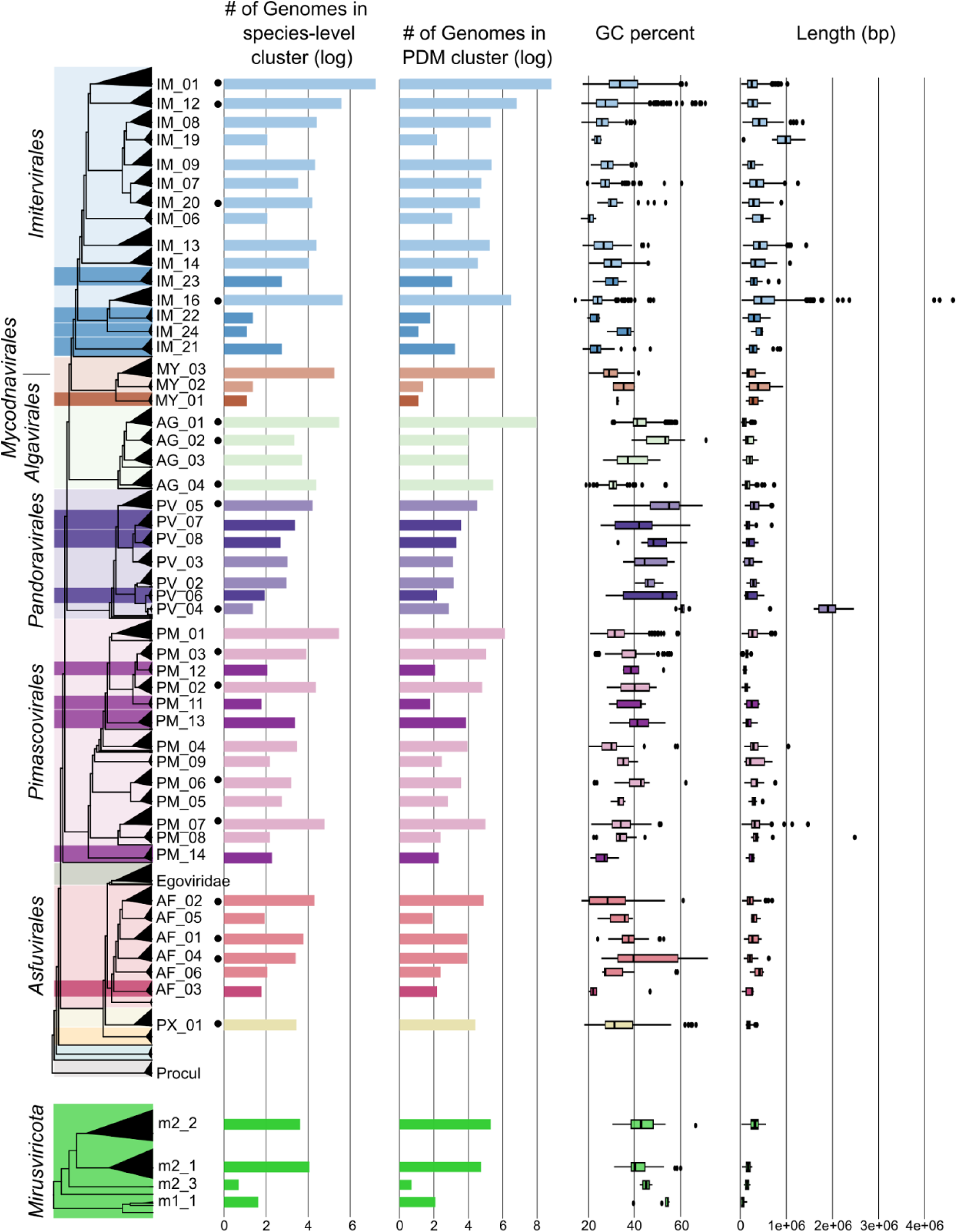
Expanded phylogenetic diversity of the *Nucleocytoviricota* and *Mirusviricota* phylums. The phylogenetic tree displays both established and newly proposed viral families within *Nucleocytoviricota*, with newly proposed viral families highlighted in darker shades within their respective *Nucleocytoviricota* orders. *Mirusviricota,* the newly proposed *Duplodnaviria* phylum, is displayed with their proposed families based on similar RED score analysis as *Nucleocytoviricota.* Families containing at least one isolate (laboratory-cultured representative) are marked with a solid black circle. For some families, four metrics are visualized: (1) logarithmically-transformed counts of genomes in sequence-based (species-level) clusters; (2) logarithmically-transformed counts of genomes in phylogenetic distance matrix (PDM) clusters; (3) genomic GC content percentage and; (3) assembly length in base pairs. Imitervirales family IM_09 separates into three families (IM_09, IM_19, IM_20) and Asfuvirales family AF_01 splits into four families (AF_01, AF_04, AF_05, AF_06). These are not considered new families, but rather a polyphyletic family split into new families following RED score values.

The GVOG7 phylogenetic tree additionally revealed a clade of *Imitervirales* genomes that split from the main clade (Figure 2), echoing findings from previous studies (Myers *et al*., 2024). This clade comprises two previously identified *Imitervirales* families: IM_18 (hereafter known as MY_03) and IM_02 (hereafter known as MY_02; *Mycodnaviridae*), and a newly identified family (hereafter known as MY_01). Their members contain similar GC percentage and assembly length as the larger *Imitervirales* group. However, its monophyly outside of *Imitervirales* is strongly supported, with a RED score consistent with the order ranking. The most robustly supported position for this clade is as a basal group to the *Imitervirales*, indicated by high support values (99/79) at the common node in the final tree. In contrast, there is lower support for an alternative topology that positions this clade as a sister group to *Algavirales*, a configuration observed in the GVOG4 and GVOG8 trees (Supplemental Figures S2 and S3).

This discrepancy is further highlighted in single gene trees (Supplemental Figure S4), where the placement varies: Packaging ATPase (GVOG760) and DNA polymerase family B (GVOG054) support positioning within *Imitervirales*, while Poxvirus Late Transcription Factor VLTF3 (GVOG890) supports the sister-to-*Algavirales* topology. Interestingly, DNA topoisomerase II (GVOG461) and Transcription initiation factor IIB (GVOG172) show a relationship where these genomes form a clade with other *Imitervirales* genomes outside the main *Imitervirales* and appear as a sister clade to *Algavirales*. One marker DEAD/SNF2-like helicase (GVOG013) unexpectedly places these genomes with *Asfuvirales*. Additionally, two markers DNA-directed RNA polymerase beta subunit (GVOG022; see Methods for additional information on GVOG022) and DNA-directed RNA polymerase alpha subunit (GVOG023) involve only a limited number of genomes, exhibiting high uncertainty due to restricted phylogenetic signals from the large number of taxa and short sequence lengths. Consequently, we designate the divergent clade as a new taxonomic order tentatively named *Mycodnavirales*, with the name derived from *Mycodnaviridae,* a proposed family of giant viruses associated with early-diverging fungi (Myers *et al*., 2024; Perini *et al*., 2024).

Furthermore, in our expanded data set we observe the division of known families into multiple non-monophyletic clades, which we denote with the clade name and a new number identifier. This phenomenon is evident in the *Imitervirales* family IM_09, which separates into three distinct families (IM_09, IM_19, IM_20), each comprising a mix of IMG/M genomes and genomes from other sources. A similar pattern is observed in the *Asfuvirales* family AF_01, where the family splits into four families (AF_01, AF_04, AF_05, AF_06). Each of these newly defined families (except AF_05) contains genomes from multiple independent sources, supporting their taxonomic validity. Second, former singleton genera now form well-supported families through the addition of related genomes. We identified four families that emerged from individual singletons (AF_03, MY_01, IM_21, IM_24) and four additional families that formed from pairs of singletons (PM_14, PV_07, PV_08, IM_23). This transformation from singleton genera to robust families demonstrates how additional genomic sampling can reveal previously hidden viral diversity and establish more accurate phylogenetic relationships.

### Expansion of the the new viral phylum *Mirusviricota*

*Mirusviricota* is a new phylum of giant viruses linked to herpesviruses (Gaïa *et al*., 2023), and seem to be associated with diverse eukaryotes, in particular protists and algae (Zhao *et al*., 2024; Archibald *et al*., 2025). We recovered 611 *Mirusviricota* MAGs from biomes spanning aquatic systems (*i.e.*, marine, freshwater, non-marine saline/alkaline, deep subsurface, and thermal springs), terrestrial systems (*i.e.,* soils and rock endoliths), plant-associated niches (*i.e.,* roots, peat moss, and phyllosphere) and engineered environments (*i.e.,* anaerobic bioreactors, soil microcosms, and plant-growth chambers). We combined our newly recovered *Mirusviricota* MAGs from the IMG/M dataset, with 88 *Mirusviricota* MAGs from TARA Oceans (Gaïa *et al*., 2023).

Dereplication of *Mirusviricota* MAGs resulted in 111 species-level-clusters and 298 singletons. Only 145 clusters and singletons contained enough markers to be further clustered with phylogenetic distance clustering (PDM), resulting in 55 clusters and singletons, with 2 clusters solely consisting of novel genomes (*i.e.,* IMG/M genomes). The complete taxonomic classification for all 55 Mirusviricota representatives follows a sequential naming convention (orders: m1/m2, families: m1_1/m2_1, genera: m2_1g_1) (Figure 2). The majority of PDM-clusters (n = 49) are positioned with *Pandoravirales*, consistent with previous phylogenetic reconstructions (Fang *et al*., 2025). Following a similar RED score-based approach as we did for *Nucleocytoviricota*, we were able to establish a taxonomic framework of *Mirusviricota* comprising two orders (Supplemental Tables 3 and 4). To better understand the evolutionary relationship of *Mirusviricota* and *Nucleocytovirictoa*, we built a combined species tree (Supplemental Figure S5). In this tree, the *Mirusviricota* m2 order contains 3 families subdivided into 13 genera (each containing multiple genomes), with one genus comprising only two genomes that was retained due to the specific characteristics of *Mirusviricota*. This genus occupies a position basal to the common node containing 47 other *Mirusviricota* sequences and a *Pandoravirales* genome (“sylv”), which likely represents long branch attraction as evidenced by low support values. The *Mirusviricota* m1 order, consisting of five genomes, appears basal to all *Megaviricetes*. While support for grouping these five genomes together is robust, their position outside the main *Mirusviricota* group remains uncertain. This clade could represent a genuine evolutionary separation through independent acquisition of *Nucleocytoviricota* genes or an artifactual placement given their relative proximity to the main group and the moderate support for internal nodes within *Megaviricetes*. This second order contains a single family with all genomes classified in independent genera. One outlier genome, clusters near the basal *Chitovirales* lineage. Nevertheless, our RED score analysis supports proposing *Mirusviricota* as its own phylum-level lineage.

### Genetic code diversity improves annotation in giant viruses

Alternative genetic codes, including stop codon reassignments, can result in incomplete annotation of protein-coding genes if unaccounted for, yet their distribution and impact in giant viruses remain largely unexplored (Shackelton and Holmes, 2008). Our analysis of 8,508 dereplicated GVMAGs showed improvements in protein-coding gene prediction in *Nucleocytoviricota* when applying alternative genetic codes (Figure 3). Leveraging alternate genetic codes, we performed a comparison of giant virus-specific gene calling optimization with GVClass and the widely used Prodigal with the -p meta option using a two-metric (Δ complete best hits vs. Δ average best-hit score) quadrant analysis (Figure 3A; Supplemental Figure S6) (Hyatt *et al*., 2010; Pitot, Brůna and Schulz, 2024). Of the 8,508 genomes analysed, 6,965 (∼82%) fall directly on one of the axes *(i.e.,* the two programs perform identically for at least one of the two metrics). The remaining 1,543 genomes show a clear directional shift with 1,066 genomes (12% of the total) residing in Quadrant I, where the optimized gene calling in GVClass recovers both more complete profile hits and higher average scores, indicating an improvement in annotation quality. The other 476 genomes (5.6%) plot in Quadrant IV, gaining additional hits at the expense of a modest decrease in score. The increase in annotation quality observed with GVClass highlights the importance of considering genetic code variations in genomic analyses, as they can lead to more accurate predictions of gene content and functionality.

**Figure 3.**
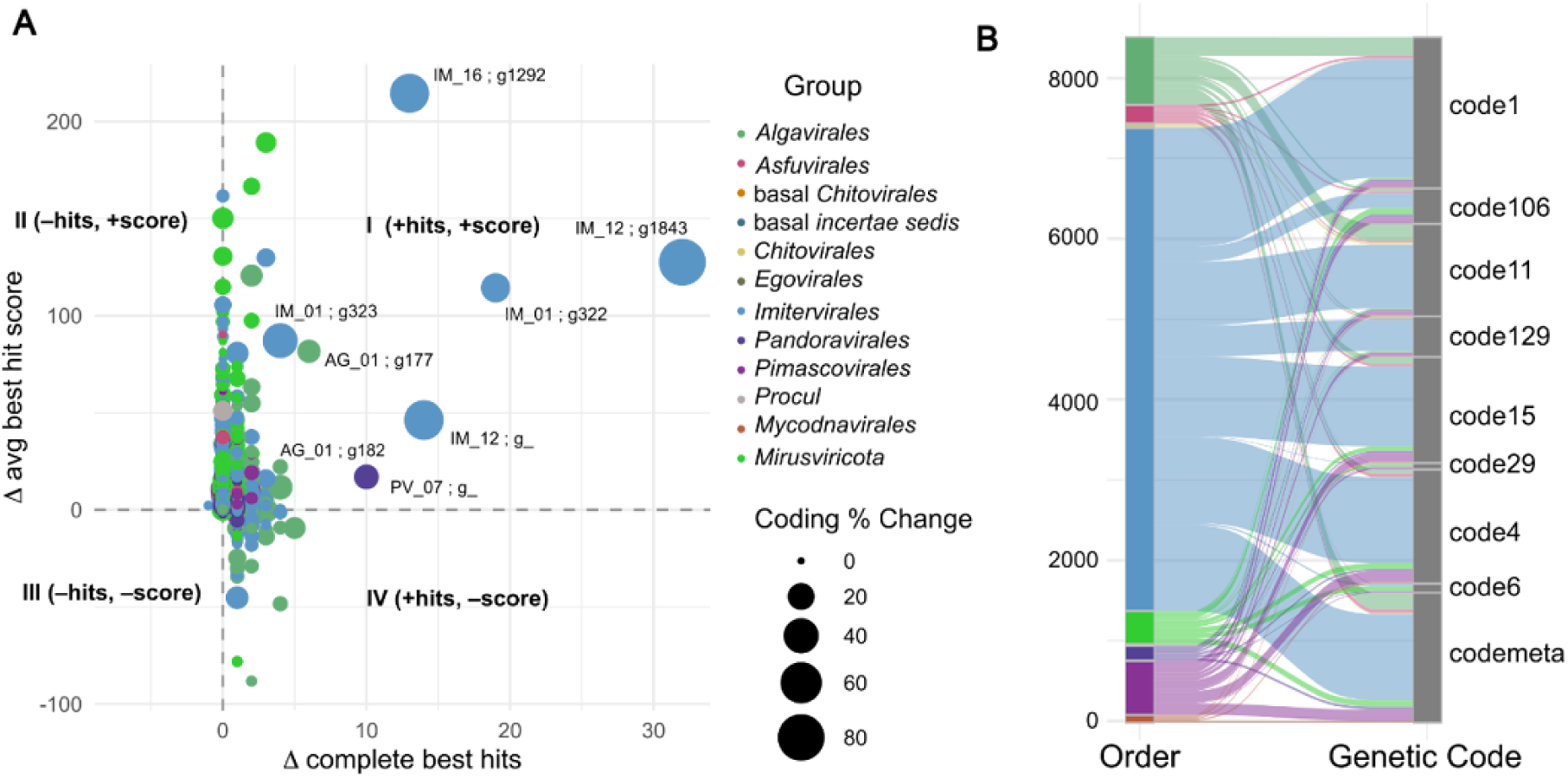
Discovery and distribution of novel genetic codes among *Nucleocytoviricota* orders and *Mirusviricota*. A) Change in complete best hits and the average best hit score between standard gene prediction (prodigal-meta) and a giant virus-specific annotation and classification tool (GVClass). Genomes with labeled family and genus names exhibit an increase of best hits of at least 3 and an increase of average best hit score of at least 15. Sizes denote the coding percent change per genome. Genomes within Quadrant 1 indicates an increase of best hits and average best hit score with GVClass, suggesting this tool is suitable for recovering previously unidentified proteins. B) Taxonomic distribution of predicted genetic codes across different orders, revealing lineage-specific patterns of genetic code usage.

To determine which specific genetic codes provided the greatest prediction accuracy across our dataset, we conducted a comparative performance analysis of multiple translation tables (Figure 3B; Supplemental Figure S7). The standard genetic code (code 1) proved to be the best optimized overall code for giant virus protein predictions, followed by one of the pre-trained models for codes 1, 4, and 11 (-p meta flag in prodigal; codemeta). Genetic codes with a single stop codon reassignments were favored by giant virus genomes more than those with multiple reassignments. For example, code 106 (reassigning TAA to glutamine) and code 129 (reassigning TAG to tyrosine) were more optimal than their counterparts, code 6 and code 29, respectively. Different viral families showed optimization for specific genetic codes. Members of *Chitovirales* showed optimized gene calling at code 11, making it the only family that showed the best optimization for a code outside of code 1 or pretrained models (-p meta flag in prodigal; codemeta). Currently, all confirmed giant virus isolates have been isolated from hosts that use the standard genetic code 1 (*e.g.,* amoebae, algae and other protists). Host-virus genetic code alignment is believed to facilitate efficient translation of viral proteins during infection (Bahir *et al*., 2009; Colson *et al*., 2013). Within a shared genetic code, hosts and viruses may still differ in their codon usage preferences. As is the case for multiple amoeba species, codon usage between giant viruses and their hosts differ, suggesting that codon usage matching alone may be a poor predictor of host range (Willemsen, Manzano-Marín and Horn, 2025). Nevertheless, considering alternative genetic codes in host prediction can reveal plausible candidates for giant viruses infecting new hosts not yet identified.

Two *Imitervirales* GVMAGs showed an increase in coding density and recovery of additional functional genes when using code 6. The first one in the family IM_12 increased its coding density from 13% to 93%, complete best hits increased from 2 to 34, and estimated genome completeness from 3 to 42%. The second one in family IM_16 showed a 52% coding density improvement, 13 additional complete best hits, and an increased genome completeness from 15% to 25%. Code 6 translates both TAA and TAG to glutamine rather than stop and is found in some species belonging to ciliates, algae and diplomonad protists (Schneider, Leible and Yang, 1989; Hoffman *et al*., 1995; Keeling and Doolittle, 1996). Currently, the only association of a giant virus to a protist that uses genetic code 6 involves the ciliate *Paramecium bursaria* that harbors a symbiotic algae (*Chlorella spp.*) in its cytoplasm, which is infected by giant viruses (Van Etten, 2003). However, there are currently no reports on giant viruses that directly infect *P. bursaria*’s nucleus or use its translation machinery, and the two genomes recovered in this study represent the first plausible candidates for giant viruses that may infect hosts using genetic code 6.

### Giant endogenous viral elements (GEVEs) are widespread across datasets

To identify putative hosts, we mined for giant endogenous viral elements (GEVEs) in publicly available eukaryote genomes. We identified a total of 150, 807, and 84 genomes with putative GEVEs from IMG/M, MycoCosm and Phycocosm, respectively, using a nearest neighbor phylogeny-based search (Grigoriev *et al*., 2014, 2021; Chen *et al*., 2019) (Supplemental Figure S8). Our analysis uncovered GEVEs in lineages where they were not previously known, further expanding the recognized host range of giant viruses. These new putative hosts of giant viruses include Glaucophyta and Rhodophyta. We detected additional GEVEs in well-studied host groups such as Symbiodiniaceae dinoflagellates (Benites, Stephens and Bhattacharya, 2022) and Phaeophyceae brown algae (Stramenopiles) (Mckeown *et al*., 2025). In green algae (Chlorophyta), where latent GEVEs have been experimentally studied (Moniruzzaman, Weinheimer, *et al*., 2020; Erazo-Garcia *et al*., 2025), we again found viral fragments.

*Nucleocytoviricota* and *Mirusviricota* GEVEs were also found in parasitic eukaryotes, including cordyceps fungi and the human pathogen *Trichomonas.* Endogenization of smaller, Maverick/Polinton-like viruses, which can parasitize giant viruses, are also widespread in *Trichomonas* (Pritham, Putliwala and Feschotte, 2007; Bellas *et al*., 2023). Overall, our findings demonstrate that integration of giant virus DNA is widespread across diverse eukaryotic lineages.

### Shared genes and enriched functions across giant virus orders

The gene repertoires of giant viruses encompass both conserved functions and diverse auxiliary genes, yet the majority of their coding potential remains uncharacterized (Abergel, Legendre and Claverie, 2015; Sun *et al*., 2020). Ortholog clustering enables a systematic assessment of this functional landscape across lineages. Comparative genomic analysis of 2,504,503 genes from dereplicated GVMAGs identified 135,998 orthologs (ProteinOrtho), spanning 1,693,149 genes. Only 75,781 orthologues were shared by at least three genomes (1,571,367 genes). Of the orthologs shared between at least three genomes, only 10,132 (14%) contained paralogs (as defined as multiple genes from a single genome in an orthogroup), indicating that the vast majority (86%) are composed of single-copy genes. As giant virus genomes are defined by a high number of gene duplications (Machado *et al*., 2023), this unexpected low frequency of orthogroups with paralogs warrants further investigation to determine if this reflects a genuine biological phenomenon or a methodological artifact related to the number of genomes analyzed. Furthermore, only 24,783 (33%) of these orthogroups had functional annotations assigned by EggNOG-mapper, highlighting a significant proportion of orthogroups of unknown function. Known functions encompass diverse biological processes, including protein synthesis and modification, and DNA damage response and repair.

Orthogroup analysis revealed distinct distributions of orthologs across the *Nucleocytoviricota* orders (Supplemental Table 6, Supplemental Figure S9). *Imitervirales* exhibited the highest number of order-specific orthologs (n = 47,541, 63% of total orthologs, 42% of total genes in orthogroups, 34% of all genes in *Imitervirales*), followed by *Pimascovirales* (n = 5,117, ∼7% of total orthologs, ∼2% of total genes in orthogroups). *Imitervirales* and *Pimascovirales* share 4,008 orthologs that are exclusive to both groups. When considering *Imitervirales* and the newly proposed *Mycodnavirales* order, 636 orthologs were shared. *Mycodnavirales* possessed 433 order-specific orthologs (with an additional 9,364 singletons). Only 68 orthogroups were shared across all five major orders (*Imitervirales*, *Pimascovirales*, *Algavirales*, *Pandoravirales*, and *Asfuvirales*).

Network analysis using a ForceAtlas2 layout (Figure 4A; Supplemental Figures S10 and S11) revealed a tendency for viruses within the same order and family to cluster together. Using the ortholog result that calculates the number of edges between two species, we calculated a within-order connectivity, defined by the number of edges shared with species within the same order divided by the total number of edges where a value closer to one indicates a stronger connection to its own order compared to its connections to other orders (Supplemental Figure S12). *Imitervirales* displayed the highest median connectivity ratio (0.91). This high connectivity is further influenced by the IM_01 family, which exhibits a median within-family connectivity of 0.79 (Supplemental Figure S13). Previous studies comparing giant viruses across diverse environments typically report lower inter-order gene conservation (Schulz *et al*., 2020), a pattern we also find in all orders outside of *Imitervirales* (all mean within-order values < 0.30). It is thought that similar eukaryotic host communities and physicochemical conditions impose convergent functional constraints (Sun and Ku, 2021). The unusually high within-order connectivity in *Imitervirales* may indicate two possibilities: high connectivity reflects the large number of *Imitervirales* genomes and the high number of *Imitervirales*-specific edges, or that, unlike other orders, their gene content is more strongly shaped by consistent host associations and environmental pressures across diverse habitats.

**Figure 4:**
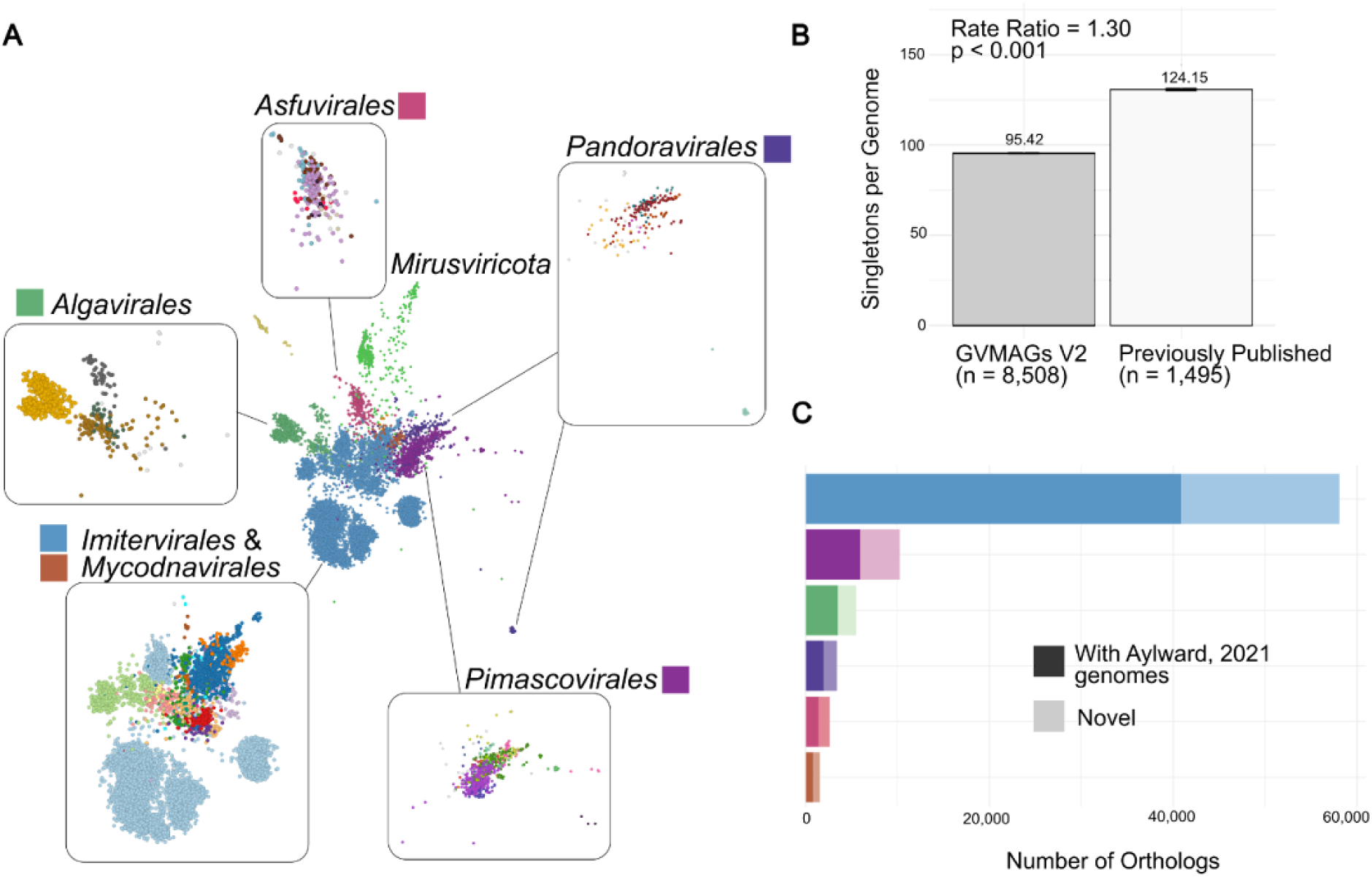
Orthologous gene families across *Nucleocytoviricota* and *Mirusviricota*. A) A bipartite network where genomes are connected through shared orthologous gene families (full network with all edges and ortholog nodes available in Supplemental Images). For clarity, edges and orthologs nodes have been removed. Network clusters colored by viral order. Inset panels: Order-specific subnetworks colored by family, revealing finer-scale patterns of gene sharing within taxonomic groups. B) Rate of singletons per dataset of our new GVMAGs V2 (n = 8,508) and a reduced dataset of only previously published giant virus genomes (n = 1,495). C) Quantification of orthologous gene families per order, with two-tone coloring indicating ortholog conservation, darker portions represent orthologs shared with genomes from the published giant virus framework (Aylward *et al*., 2021), while lighter portions show newly identified orthologs with no members from this framework.

A total of 811,354 giant virus genes represented singletons, genes not assigned to any orthogroup. This accounts for 32.4% of all total genes analyzed in this study. To assess how genome sampling affects orthogroup formation, we compared singleton rates (Figure 4B) before and after adding 7,013 genomes (those recovered from the IMG/M dataset not included in previous studies) to previously published 1,495 representatives (includes genomes from TARA oceans, Aylward *et al.,* 2021, isolate genomes and deep sea sediment genomes). Using a Poisson rate test, we found that the singleton rate significantly decreased from 124.15 to 95.4 singletons per genome (rate ratio = 1.30, p < 2.2×10⁻¹⁶) after adding the new genomes. This 1.3-fold decrease in singleton genes demonstrates that expanding genomic sampling strongly reduces the number of orphan genes by revealing previously hidden orthologous relationships. These results suggest that the current number of apparent singletons in giant virus genomes is likely inflated due to incomplete sampling of genomic diversity, and that many genes currently classified as taxon-specific may actually have undiscovered homologs in yet-to-be-sequenced organisms. In addition to reducing the singleton rate, our expanded dataset also led to a significant increase in the number of identified orthologs (Figure 4C). We identified 32,551 orthologs (out of the total 75,781 orthologues shared by at least three genomes) that do not contain genomes reported in the previous framework (Aylward *et al*., 2021). This increase demonstrates the importance of comprehensive genomic sampling with newly sequenced genomes to understand the true extent of giant virus diversity and functional potential.

Having established patterns of gene sharing and taxonomic connectivity, we investigated differences in functional capacity using completeness of metabolic modules. Analysis of metabolic module completeness revealed variations across giant virus families (Figure 5A and Supplemental Figure S14). To identify family-specific enrichment of metabolic modules, we binarised module completeness scores (present/absent), compared each family’s module frequency against that of all other families with 2 × 2 contingency tables, applied Fisher’s exact tests, and adjusted the resulting P-values using the Benjamini-Hochberg procedure. We identified 220 distinct modules enriched across distinct families (Supplemental Figure S15). For example, *Algavirales* showed enrichment for genes involved in the formaldehyde assimilation ribulose monophosphate pathway (M00345), which includes a sugar manipulation enzyme previously identified in prasinoviruses (Weynberg, Allen and Wilson, 2017). In contrast, *Imitervirales* genomes harbor genes for a different sugar manipulation process, the formaldehyde assimilation xylulose monophosphate pathway (M00344), as shown in Ha, Moniruzzaman and Aylward, 2021.

**Figure 5.**
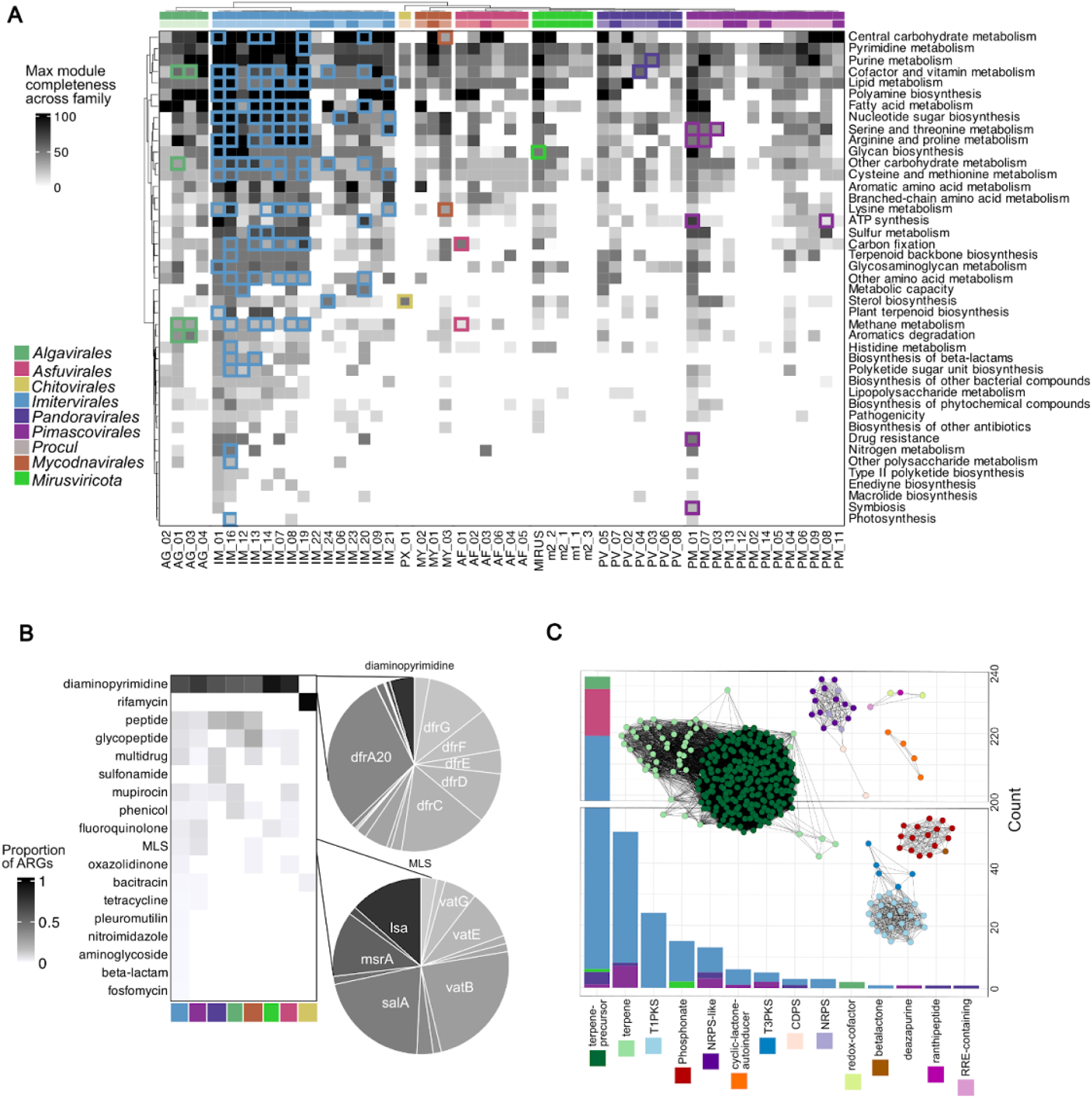
Functional capabilities across the *Nucleocytoviricota* and *Mirusviricota*. A) Overview of max completeness of various metabolic modules across different viral families. Hierarchical clustering was applied to both axes, however, clustering for families was done within order. Darker colors in the orders indicate newly identified families from this paper. Tiles with colored borders indicate gene categories that contain order-specific modules (extended figure can be found in Supplemental Figure S15). B) Antibiotic resistance genes (ARGs) within each *Nucleocytoviricota* order and *Mirusviricota*. Proportion of ARGs correspond to the proportion of each ARG category within each order. Two gene categories were expanded to show the proportion of genes present in the gene category. C) Total count of biosynthetic gene clusters (BGCs) in each BGC category. Bar chart colors correspond to the order colors shown in (A). Network plots generated using the Cosmos layout in cytoscape are colored by the BGC category where the corresponding color is next to labels. Abergel, C. *et al*. (2007) “Virus-Encoded Aminoacyl-tRNA Synthetases: Structural and Functional Characterization of Mimivirus TyrRS and MetRS,” *Journal of Virology*, 81(22), pp. 12406–12417. Available at: https://doi.org/10.1128/JVI.01107-07.

We additionally identified enriched modules that open new avenues for metabolic discovery. In particular, *Algavirales* is significantly enriched in three modules for aromatic degradation (M00545, M00551, M00569). These modules are found in *Algavirales* genomes recovered from marine and freshwater environments; including shale gas reservoirs, estuarine waters, lakes, rivers, ponds, and salt marshes. Giant viruses are already known to encode diverse metabolic genes that modulate host carbon metabolism during infection (Moniruzzaman, Martinez-Gutierrez, *et al*., 2020); the presence of aromatic degradation genes in *Algavirales* likely reflects a similar strategy to infect host cells and reprogram host metabolism. The same meta-cleavage modules found in *Algavirales* support the degradation of natural and industrial pollutants in other microbial species, and could be potentially used in bioremediation (Ferrández, Garciá and Díaz, 1997; Elbehery and Deng, 2022; Ru *et al*., 2023; Zhu *et al*., 2024). Thus, giant viruses may represent an overlooked reservoir, and vehicle, of catabolic genes with potential for biotechnological exploitation.

### *Nucleocytoviricota* and *Mirusviricota* as reservoirs for biotechnological potential

One important category often discussed for biotechnological exploration are antibiotic resistance genes (ARGs) and antimicrobial biosynthetic gene clusters. Recent work found that giant viruses can act as reservoirs of ARGs, with 12 ARG types being encoded by giant viruses (Yi *et al*., 2024). Our genome dataset expanded this work (Figure 5B, Supplemental Table 7).

Consistent with previous work, trimethoprim resistance genes (*dfr*) are the most abundant ARGs in our dataset (n = 1,604 genes). Similarly, we found a large number of glycopeptides (*van;* n = 144 genes) and mupirocin resistance genes (n = 114). Our dataset also showed evidence of genes related to beta-lactam (two genes) and nitroimidazole (five genes) resistance that were absent from a recent larger scale survey of ARGs across giant virus genomes (Yi *et al*. 2024). This survey leveraged the previous giant virus taxonomy framework (Aylward et al., 2021), which included 1,382 species-level representatives compared to the 8,508 analyzed here.

Beta-lactamase genes have been reported in giant viruses (Colson et al., 2020; Rigou et al., 2022), indicating that the earlier framework missed known functions likely due to limited genome representation at the time. Similarly, nitroimidazole resistance genes have been previously reported in giant viruses, but not identified in Yi *et al*., 2024 (Chatterjee and Kondabagil, 2019; Li *et al*., 2024). The beta-lactam and nitroimidazole resistance genes are found in three different ecological niches: freshwater lake, estuary water, and saline lake water. Beta-lactams are among the most widely prescribed antibiotics (Yimenu *et al*., 2019; Centers for Disease Control and Prevention, 2022), and nitroimidazoles are often used to treat anaerobic bacterial and protozoal infections (Alexander, Gielen and Sartorelli, 1986). By recovering previously known but overlooked genes, absent from recent large scale surveys, our work provides an updated view of ARGs within giant viruses and their presence in diverse aquatic ecosystems highlights the need to consider giant viruses in antimicrobial resistance surveillance frameworks. Giant viruses may serve as reservoirs of biotechnologically relevant functions, as understanding their resistance mechanisms could enable the development of new genetic tools to address antibiotic resistance.

To further explore the functional potential of giant viruses, we surveyed biosynthetic gene clusters (BGCs) across our curated database using antiSMASH (Figure 5C, Supplemental Table 8) (Blin *et al*., 2025). We identified BGCs in 326 giant virus genomes, highlighting secondary metabolic capacities within *Nucleocytoviricota* and *Mirusviricota*. The majority of predicted clusters fell within terpene-precursor (n = 233) and terpene (n = 49) categories. While terpene biosynthesis has only been examined in four viral genomes to date (Jung *et al*., 2023; Park *et al*., 2025), our data reveal that putative terpene biosynthetic genes are far more widespread, spanning all major *Nucleocytoviricota* orders, including *Algavirales, Imitervirales, Asfuvirales, Pandoravirales,* and *Pimascovirales*, as well as the new phylum, *Mirusviricota*. This suggests that the capacity for terpenoid production, which are compounds involved in membrane modification, signaling, or defense (McGarvey and Croteau, 1995), may be a common and ecologically relevant feature of giant viruses, with potential implications for virus-host interactions. Beyond terpenes, our analysis uncovered BGCs associated with polyketides (T1PKS and T3PKS) and peptide-derived compounds (nonribosomal peptide synthetases (NRPS)-like), categories well-known for their antimicrobial activities (O’Hagan, 1991; Cane, Walsh and Khosla, 1998; Cane and Walsh, 1999). These gene clusters were primarily concentrated within *Imitervirales*, *Pandoravirales*, and *Pimascovirales*, raising the possibility that some giant viruses may influence microbial community dynamics within host niches. We also observed order-specific patterns, including redox-cofactor clusters uniquely in *Algavirales*, and BGCs such as RiPP recognition element (RRE)-containing and ranthipeptide clusters exclusively in *Pandoravirales*, hinting at unexplored diversity in viral small-molecule biosynthesis. Likewise, the detection of deazapurine, a nucleotide modification pathway (Hutinet, Swarjo and de Crécy-Lagard, 2017), in *Pimascovirales*, phosphonate gene cluster in *Imitervirales* and *Mirusviricota*, and quorum-sensing-like cyclic-lactone autoinducer clusters in both *Imitervirales* and *Pimascovirales*, underscores the functional complexity of giant virus genomes.

## CONCLUSION

Our study significantly expands the known diversity and functional potential of *Nucleocytoviricota* and *Mirusviricota*. Through phylogenomic and comparative genome analyses, we identified new viral diversity, characterized both conserved and lineage-specific gene content, and revealed novel genetic code usage and metabolic potential across major viral clades. Building upon previous classifications (Aylward et al., 2021), we identified one new order, 13 new families and 712 novel genera, including numerous singleton genomes that likely represent yet-undescribed lineages. These findings highlight the current underestimation of giant virus diversity and the need for further genome-resolved metagenomics, especially in underexplored habitats such as polar regions and the deep sea.

While most genomes are best annotated with the standard genetic code, several showed notable improvements in gene prediction using alternative genetic code. These cases, which were checked for contamination, raise the possibility of previously unrecognized host-virus associations and suggest that the landscape of giant virus-host coevolution may be broader than currently appreciated. Our survey of publicly available eukaryotic genomes further expands the known host range of giant viruses through discovery of GEVEs into algae, several parasitic fungi, and the human pathogen *Trichomonas*. This indicates that endogenization of giant virus DNA is a recurring feature of eukaryotic genome evolution.

Despite our study expanding the coding potential of giant viruses, the vast majority of genes (∼67%) remain uncharacterized, reflecting both the novelty of giant virus gene content and the limitations of current annotation databases. Several limitations should be considered.

Incomplete metagenome-assembled genomes may bias our view toward more readily assembled viral genomes and obscure the full spectrum of viral diversity. Uneven environmental sampling leads to overrepresentation of certain ecosystems and host types. Limited host information constrains ecological interpretations and understanding of genetic code variation. Nevertheless, even within these constraints, our large-scale genome mining uncovered an expanded repertoire of biosynthetic gene clusters and antibiotic resistance genes, positioning giant viruses as an underexplored reservoir of novel enzymes, metabolic pathways, and bioactive compounds with potential applications in biotechnology, synthetic biology, and drug discovery. Thus, our findings provide a foundation for future experimental efforts aimed at exploring the biotechnological potential of giant virus encoded genes and pathways.

## METHODS

### Dereplication & phylogenetic inference of GVMAGs

A total of 18,727 genomes were downloaded from multiple sources, including 14,914 MAGs recovered from the IMG/M database (Villada *et al*., 2025), 1,516 genomes from a coassembly of MAGs from the freshwater Lake Mendota (Oliver *et al*., 2024), 1,353 genomes from the 2021 giant virus taxonomy framework (Aylward *et al*., 2021), 630 genomes for the TARA Oceans sequencing efforts (Tully *et al*., 2017; Gaïa *et al*., 2023), 227 isolate giant virus genomes (Benson *et al*., 2012), 65 from a sequencing effort for protists single-cell assembly genomes (Schulz *et al*., 2024) and 22 genomes from deep sea sediments (Bäckström *et al*., 2019).

First, we did an initial dereplication using nucleotide identity. We calculated the pairwise average nucleotide identity between MAGs using skani v 0.2.1 (settings: -slow -min-af 30 -m 200) (Shaw and Yu, 2023). MAGs with at least 95% ANI and 30% alignment were considered in the same cluster (hereafter denoted as “species-level cluster”). Each species-level cluster was chosen to have one MAG representative, preferring the MAG with the highest number of the nine giant virus orthologous groups (GVOGs) markers (Aylward *et al*., 2021). Using the species-level cluster representatives and all genomes that do not fall into a species-level cluster (hereafter identified as singletons), we used sgtree (commit 2e7f93d https://github.com/NeLLi-team/sgtree) to perform phylogenetic tree construction using the GVOG7 markers (Aylward *et al*., 2021).

Briefly, sgtree aligns sequences using MAFFT v7.520, trims alignments with trimAl v1.4.rev1 (settings: -gt 0.1), and constructs the tree using Fasttree v2.1.11 (model: LG+F+I+G4) (Katoh *et al*., 2002; Capella-Gutiérrez, Silla-Martínez and Gabaldón, 2009; Price, Dehal and Arkin, 2010). We filtered out gene markers shorter than 50% of the marker median length, and then genomes with less than three GVOG4 (“GVOGm0461”, “GVOGm0022”, “GVOGm0023”, “GVOGm0054”) or two GVOG4 and three GVOG8 (“GVOGm0013”, “GVOGm0022”, “GVOGm0023”, “GVOGm0054”, “GVOGm0172”, “GVOGm0461”, “GVOGm0760”, “GVOGm0890”) markers were removed from further dereplication. To further dereplicate genomes, we used the sgtree phylogenetic tree and converted it into a pairwise distance matrix then further converted the distance into a similarity value (maximum 1) using PhyloDM (https://github.com/aaronmussig/PhyloDM). We generated a set of similarity tables, with different cutoff values for similarity, keeping the distance for similarity higher than the cutoff, from 0.98 to 0.05 with steps of 0.01. All similarity tables were clustered using MCL (inflation 2), obtaining clusters of phylogenetically close MAGs (hereafter identified as “PDM-cluster”). Using the taxonomic assignments for the genomes from the giant virus taxonomy framework (Aylward *et al*., 2021), we counted how many genomes from different genera ended in the same MCL cluster for each cutoff value. We selected the lowest cutoff number that did not collapse different genera in the same dataset. For each PDM-cluster we selected a representative genome, favoring the genomes with the highest number of OGs markers. The final genome phylogenetic tree consists of the PDM-cluster representatives, all unclustered genomes (*i.e*., singletons), and the addition of 4 Procul and 192 Egovirales genomes, resulting in 2,037 genomes. Our final tree was inferred using IQtree v2.3.5 Q.pfam+R10+F with 1000 ultrafast bootstraps and 1000 SH-aLRT in sgtree and the GVOG7 HMM markers. The obtained tree was rooted on the Proculviricetes. We additionally tested our procedure with the genomes used in Aylward et. al (2021) to see if we were able to replicate the tree. The topology of the resulting trees was dependent more on the selection of the gene markers than to the parameters of the phylogenetic inference: our tree built without correction for the incomplete lineage sorting resulted in the splitting of a part of Imitervirales, with a common ancestor with Algavirales and Pandoravirales.

To build Supplemental trees in order to consider alternative topologies, we used the PDM-clustered representatives to build trees using concatenated GVOGs and single GVOG trees. We specifically used a concatenation of four markers (GVOG4) and a concatenation of eight markers (GVOG8). For GVOG4, we kept all representatives with at least 3 GVOG4 markers, and re-ran sgtree, obtaining 1810 genomes, plus the 4 Procul and 192 Egovirales. The final tree was inferred using IQtree v2.3.5 Q.pfam+R10+F with 1000 ultrafast bootstraps and 1000 SH-aLRT. The GVOG4 tree had limited phylogenetic signal, and reduced the number of genomes usable in the phylogeny with enough markers. The relations between the orders were not clearly solved as their internal nodes often had limited support. For example, the basal taxa (*i.e.,* basal Chitovirales and basal Pokkesviricetes) were nested inside Imitervirales and inside Algavirales, respectively.

For the GVOG8 tree, to the original GVOG7 markers. First, we identified GVOG0022 sequences using HMMER v3.4 on the PDM-cluster representatives, aligned the protein sequences with MAFFT v7.520, and trimmed with trimAl v1.4.rev1 (settings: -gt 0.1). We built a single gene tree on the GVOG0022 sequences with IQtree v2.3.5 LG+F+R4. The tree was manually inspected to identify if paralogs were present, and we found two clades of Imitervirales (n = 631) showing presence of paralogs that form sister clades to Pimascovirales (Supplemental Figure S16). Therefore, we subsetted the GVOG0022 alignment, keeping the sequence with the best HHMER bitscore for each genome. The sequences were realigned and trimmed, and added to the GVOG8 concatenate, and the tree was inferred with IQtree v2.3.5 Q.pfam+R10+F with 1000 ultrafast bootstraps and 1000 SH-aLRT. The GVOG8 tree had low support for the internal nodes separating the orders, with Algavirales paraphyletic with some of its clades monophyletic with Pandoravirales, a topology inconsistent with other trees and literature.

### Taxonomic assignments of GVMAGs

To assign the taxonomic rank of our final dataset, we used the R package Castor v1.8.2 to calculate the Relative Evolutionary Distance (RED) scores for the rooted phylogenetic tree where each node has a score calculating the relative position between the root and the tips (Louca and Doebeli, 2018; Parks *et al*., 2018). The calculated RED scores were used to assign the taxonomic classification without overlaps between different levels of taxonomic ranks, using the genomes from the Framework dataset as a reference. We assigned family taxonomy if there were at least 3 genomes in the taxa, otherwise family taxonomy was classified as “*incertae sedis*”. We further assigned genus taxonomy using the calculated RED scores.

### Coding potential and inferred functional capabilities of GVMAGs

To consider alternative genetic codes for giant virus genomes, we used GVClass v1.0 (Pitot, Brůna and Schulz, 2024). GVClass tests different genetic codes and evaluates results based on a hmmsearch using the general models. Currently, GVClass will select the optimal code based on the genetic code that yields the largest number of matches to general models (*i.e.,* specific gene orthogroups) with the highest average bitscore and the highest coding density. Genome completeness is calculated as the count of genes conserved in 50% of genomes of the respective *Nucleocytoviricota* order. When needed to confirm the lack of contamination, a BLASTP with diamond v2.1.9 search was performed (Camacho *et al*., 2009; Buchfink, Reuter and Drost, 2021).

We further identified orthologous genes we used Proteinortho v6.3.3 (settings: -p=diamond -e=1e-5 -identity=25 -cov=50 -sim=0.80, --core) with all genomes included in the phylogenetic tree, or were clustered through species-level or PDM (Klemm, Stadler and Lechner, 2023). We performed a smaller ortholog search using the non-redundant GVMAGs. We used gephi to visualize the orthogroups shared by at least 3 genomes, using a bipartite graph where genomes and orthogroups are nodes and the presence of genes as weightless edges (Bastian, Heymann and Jacomy, 2009). We used a ForceAtlas2 layout with default settings and 100 iterations (Jacomy *et al*., 2014). ForceAtlas2 repels nodes from one another, while using edges to pull connected nodes together, thus creating a graph where highly connected nodes cluster together.

Finally, we assessed the completeness of metabolic modules based on the metabolic module definitions of KEGG (Kanehisa and Goto, 2000). The KEGG KOs functional annotations of every genome were extracted from the results of EggNOG-mapper v2.1.10 (Cantalapiedra *et al*., 2021); when more than one KO was assigned to a protein, all the KOs were unnested and considered individual KOs for downstream analyses. The list of KOs per genome was then passed to the script ko_mapper.py from MicrobeAnnotator v2.0.5 to map KOs to their respective modules and compute metabolic module completeness per genome (Ruiz-Perez, Conrad and Konstantinidis, 2021).

To identify family-specific enrichment of metabolic modules, we first converted module completeness scores to a simple binary variable (present/absent). Any value over 0 was considered present. For every module we then counted, within each focal family, how many genomes contained the module and how many did not, and contrasted these numbers with the corresponding counts from all other families combined. The resulting 2 × 2 contingency tables were analysed with Fisher’s exact test to obtain odds ratios and P-values, which were subsequently adjusted for multiple testing with the Benjamini-Hochberg procedure. Infinite odds ratios arising from zero counts were truncated to 5 × 10³ for downstream display.

To identify antibiotic resistance genes, we used DeepARG v1.0.2 which identifies a broader diversity of ARGs (--model LS --min-prob 0.8 --arg-alignment-identity 25 --arg-alignment-evalue 1e-10 --arg-num-alignments-per-entry 1000). For identification of biosynthetic gene clusters, we used antiSMASH 8.0 and BLASTP (-evalue 1e-5) against the Minimum Information about a Biosynthetic Gene cluster (MIBiG) database (Camacho *et al*., 2009; Blin *et al*., 2025; Zdouc *et al*., 2025). Additionally, we used BLASTP against the NR database on all contigs containing putative biosynthetic gene clusters to confirm they are nested within *Nucleocytoviricota* genes. We used BiG-SLiCE v2 (Kautsar *et al*., 2021) to cluster or BGCs by similarity and cytoscape web 1.0 (Ono *et al*., 2025) to visualize networks.

### Identification of giant endogenous viral elements

Putative endogenous viral elements (EVEs) were identified from 194 algal genomes and 1,599 eukaryotic bins in the PhycoCosm and MycoCosm (Grigoriev *et al*., 2014, 2021), along with 751 eukaryotic MAGs in the IMG/M databases with GVClass v1.0 (Pitot, Brůna and Schulz, 2024). We identified putative EVEs using a nearest neighbor search from the GVClass output where EVEs that were in proximity with an *Nucleocytoviricota* gene were used for further analysis. The putative EVEs were further assessed to check if the detected viral signature was integrated into a eukaryotic contig to avoid analyzing viruses that were not truly endogenized. To do this, we selected contigs with hits to eukaryote Benchmarking Universal Single-Copy Orthologs (BUSCO) (Manni *et al*., 2021) and Giant Virus Orthologous Groups (GVOGs) or Mirusvirus Orthologous Groups (mOGs) (Zhao *et al*., 2024) profile hidden Markov models HMMER v.3.4 (Eddy, 2011) for downstream analysis. The hosts of these high confidence EVEs were assigned taxonomy through a consensus of EukCC v2.1.3 (Saary, Mitchell and Finn, 2020) and BLASTp (Camacho *et al*., 2009) of the BUSCO hits.

## Data Availability

The complete dataset is freely available at https://portal.nersc.gov/cfs/nelli/gvmagsV2/. Additional data is available at https://yvasquez-lbl.github.io/GVMAGsV2/.

## Acknowledgements

The work conducted by the U.S. Department of Energy Joint Genome Institute (JGI) (https://ror.org/04xm1d337), a DOE Office of Science User Facility, is supported by the Office of Science of the U.S. Department of Energy operated under Contract No. DE-AC02-05CH11231. We thank all JGI users whose data has been included in this work. This project was funded by the European Community’s H2020 Programme H2020-MSCA-RISE 2019 under grant agreement No. 872767.

