## Supplemental Figures for "Genome-resolved expansion of *Nucleocytoviricota* and *Mirusviricota* reveals new diversity, functional potential, and biotechnological applications"

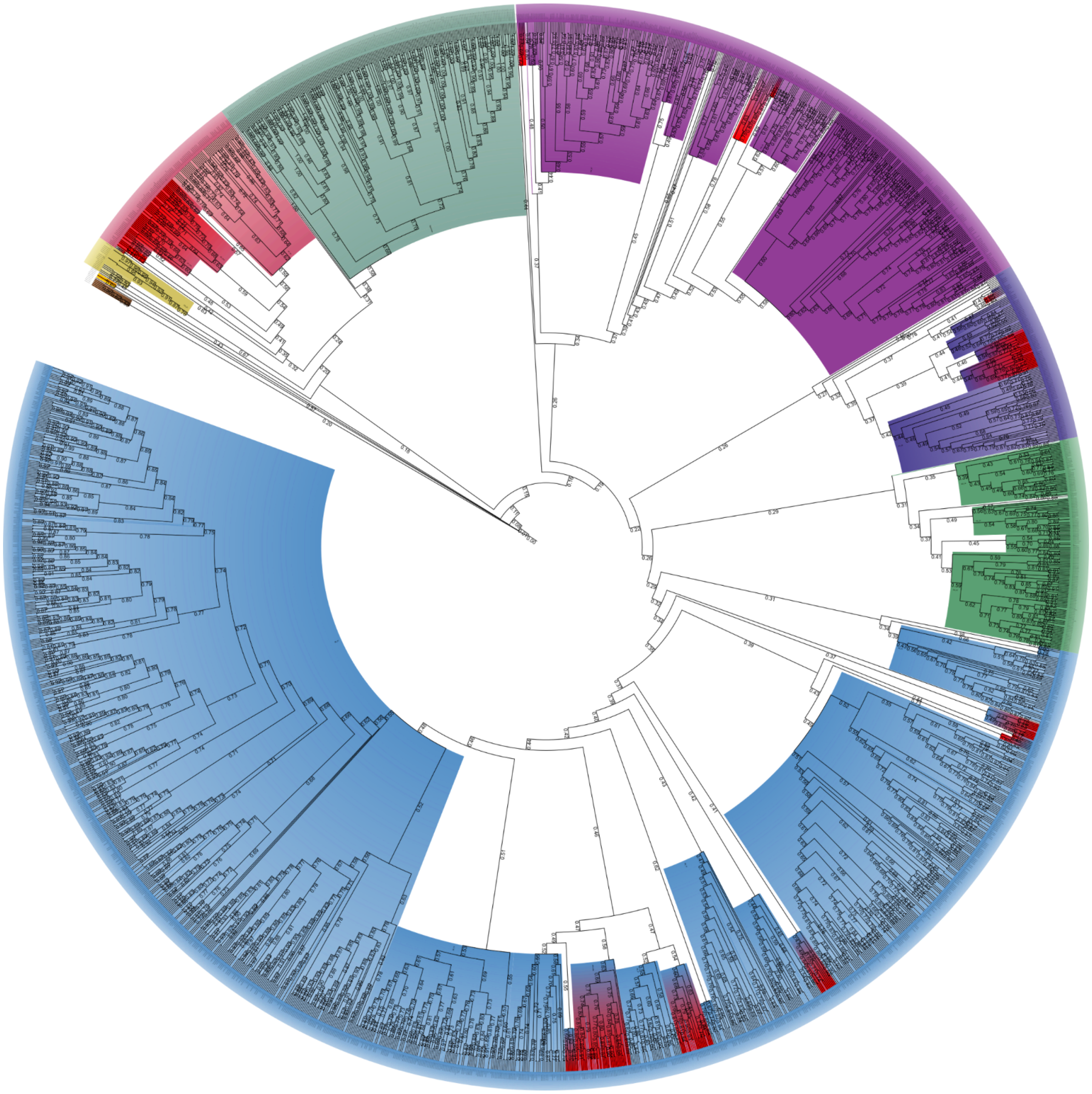

Supplemental Figure S1: Phylogenetic tree using GVOG7 markers with RED scores and taxonomic classification. The circular phylogenetic tree was constructed using seven GVOG marker genes. The tree displays distinct clades color-coded by taxonomic order. RED (Relative Evolutionary Divergence) scores are displayed on branches. Putative new families are highlighted in red throughout the tree. The interactive version of this tree is available at https://itol.embl.de/shared/fmschulz.

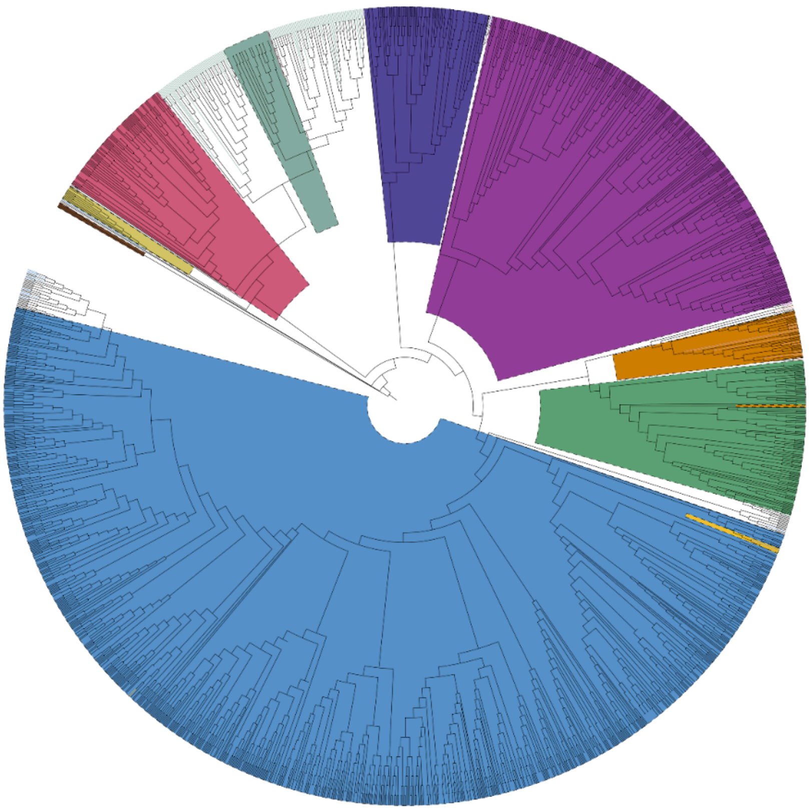

Supplementary Figure S2: Phylogenetic tree using GVOG4 markers and taxonomic classification. The phylogenetic tree was constructed using four GVOG marker genes. The tree displays distinct clades color-coded by taxonomic order. The gold-brown clade displays the split-Imitervirales genomes sister-clades with Algavirales.

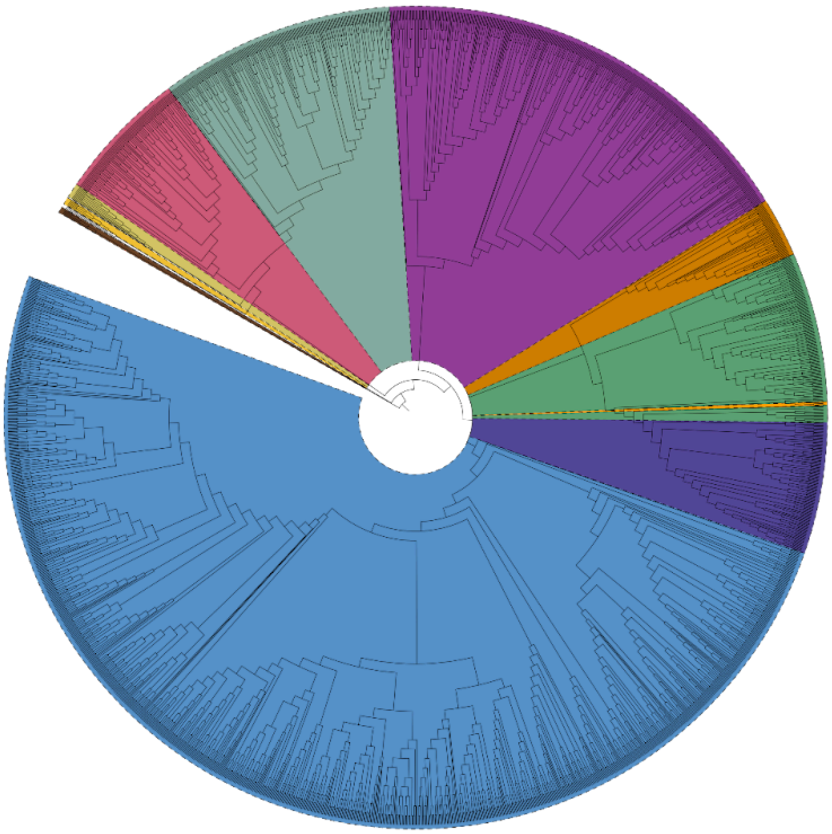

Supplementary Figure S3: Phylogenetic tree using GVOG8 markers and taxonomic classification. The phylogenetic tree was constructed using eight GVOG marker genes. The tree displays distinct clades color-coded by taxonomic order. The gold-brown clade displays the split-Imitervirales genomes sister-clades with Algavirales.

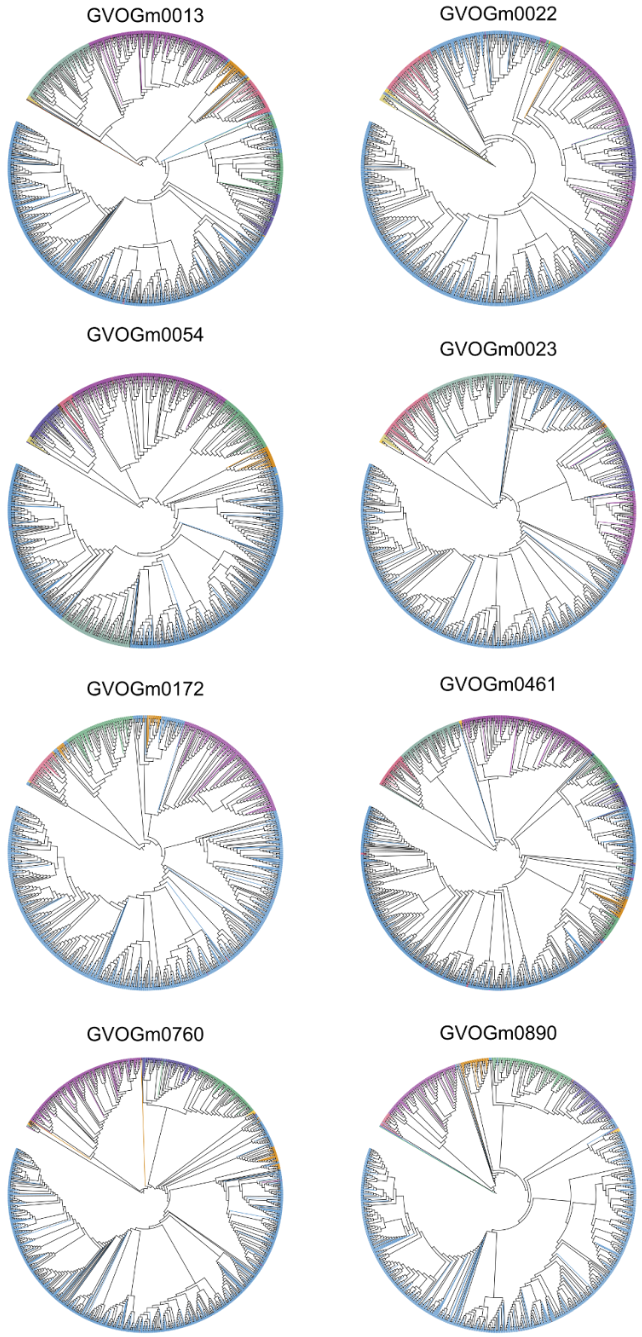


Supplementary Figure S4: Phylogenetic trees using single GVOG markers and taxonomic classification. Each phylogenetic tree was constructed using a single GVOG marker gene. The tree displays distinct clades color-coded by taxonomic order. The gold-brown clade displays the *Mycodnavirales* genomes and its inconsistent placing. The interactive version of these trees are available at https://itol.embl.de/shared/fmschulz.

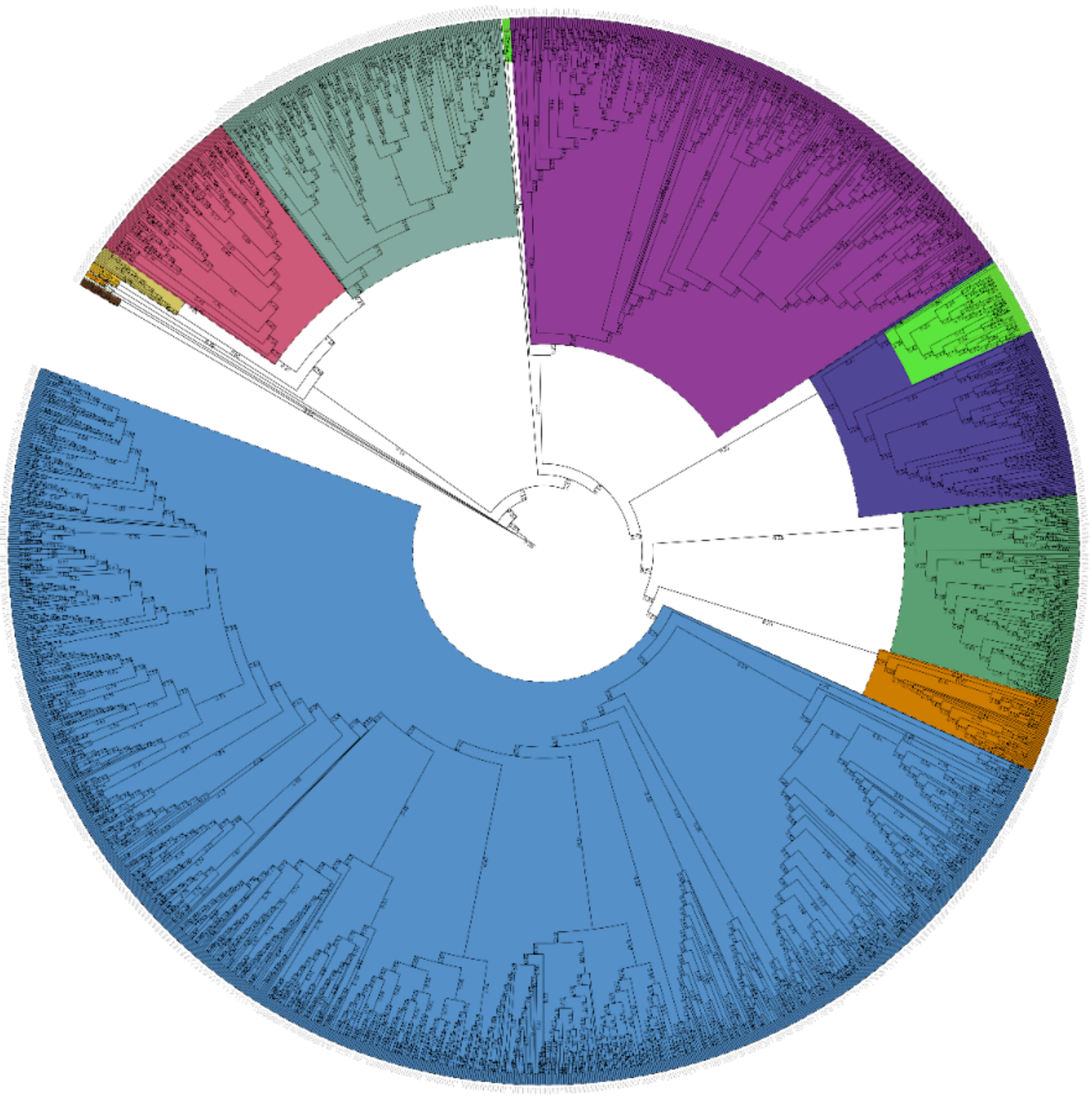

Supplementary Figure S5: Phylogenetic tree with RED scores and taxonomic classification. The tree displays distinct clades color-coded by taxonomic order. RED (Relative Evolutionary Divergence) scores are displayed on branches. The interactive version of this tree is available at https://itol.embl.de/shared/fmschulz.‘Mirusviricora’ represents the lime green clade nestled with Pandoravirales.

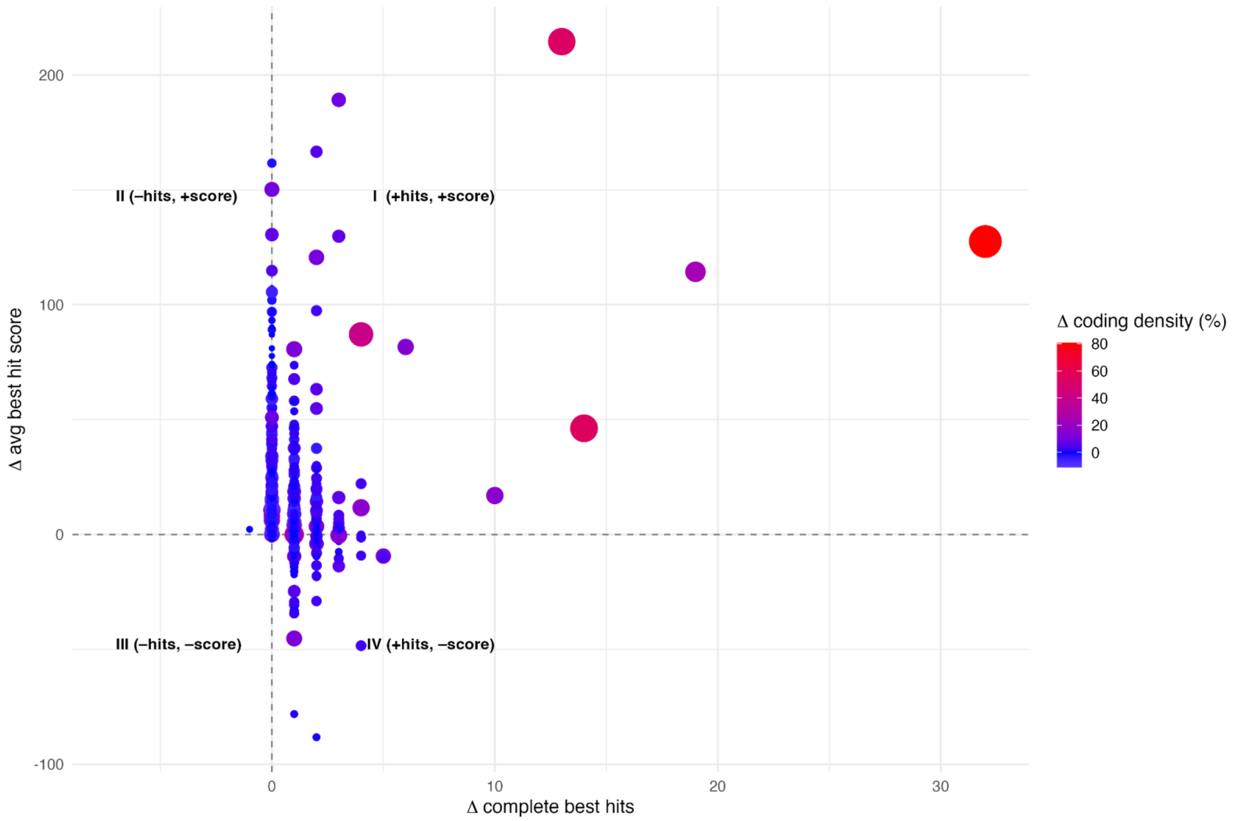


Supplementary Figure S6: Comparison of GVclass and Prodigal gene prediction performance across genomes. Plot compares the performance of GVclass versus Prodigal gene prediction, using the change in complete best hits count (gvclass - prodigal) and the change in average best hit score (gvclass - prodigal). Size and coloring correspond to the change in coding percentage (coding density) of gvclass vs. prodigal. Quadrant I (+hits, +score) represents genomes where GVclass identified more complete hits with higher scores than Prodigal; Quadrant II (-hits, +score) shows cases where GVclass found fewer hits but with improved scores; Quadrant III (-hits, -score) indicates reduced performance in both metrics; and Quadrant IV (+hits, -score) represents increased hit numbers but with lower scores. The majority of data points cluster in Quadrant II, suggesting that GVclass generally produces fewer but higher-quality gene predictions compared to Prodigal. Genetic codes in GVclass are ranked based on: complete best hits (the highest number of complete profile hits with >60% of model coverage) and the average best hit score (the average of bitscores corresponding to the best profile hits for each predicted protein).

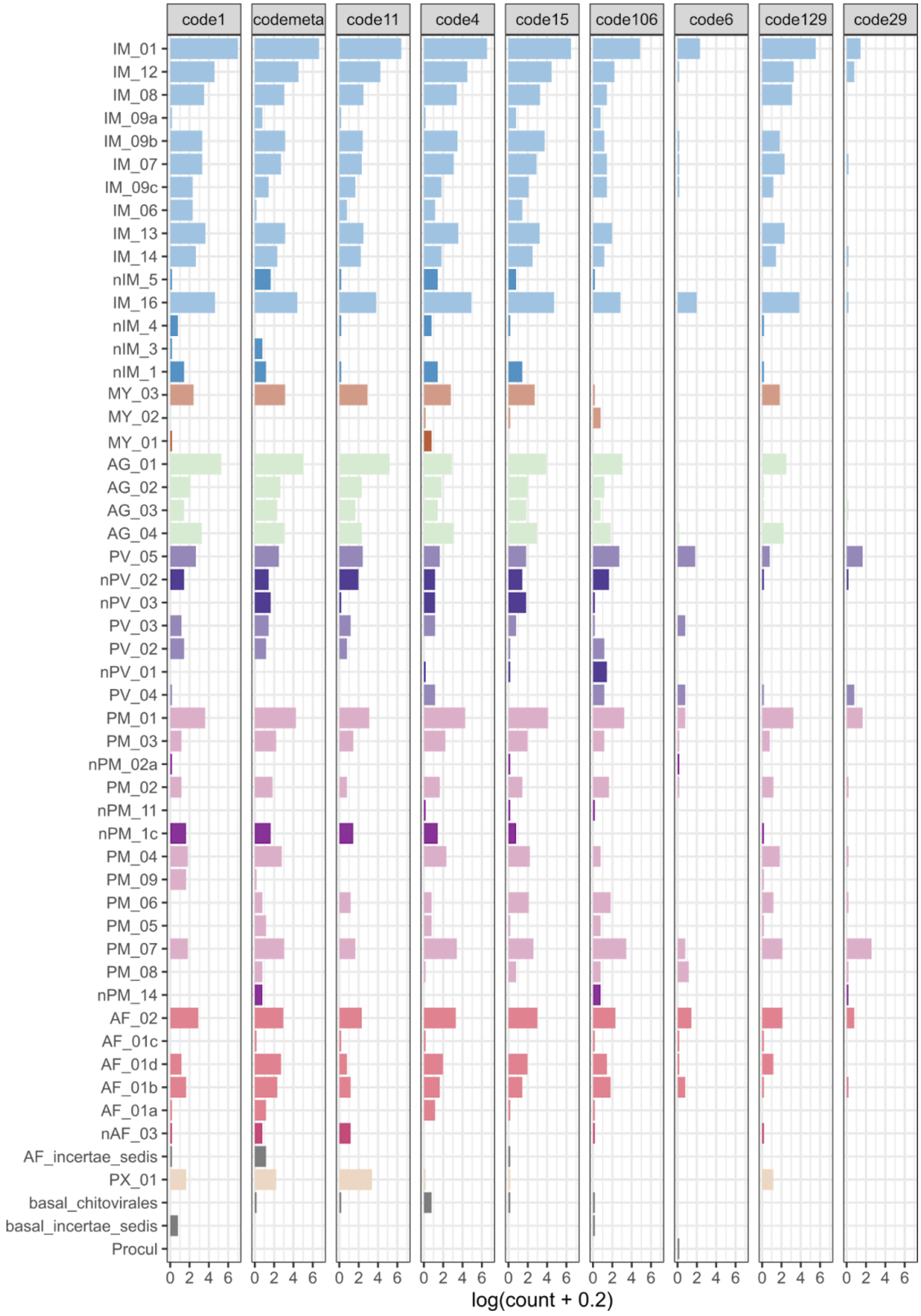


Supplementary Figure S7: Distribution of genetic codes across families. Frequency distribution of different genetic codes (code1, codemeta, code11, code4, code15, code106, code6, code129, and code29) across various families, the x-axis showing log-transformed genome counts, and families color-coded by taxonomic order.

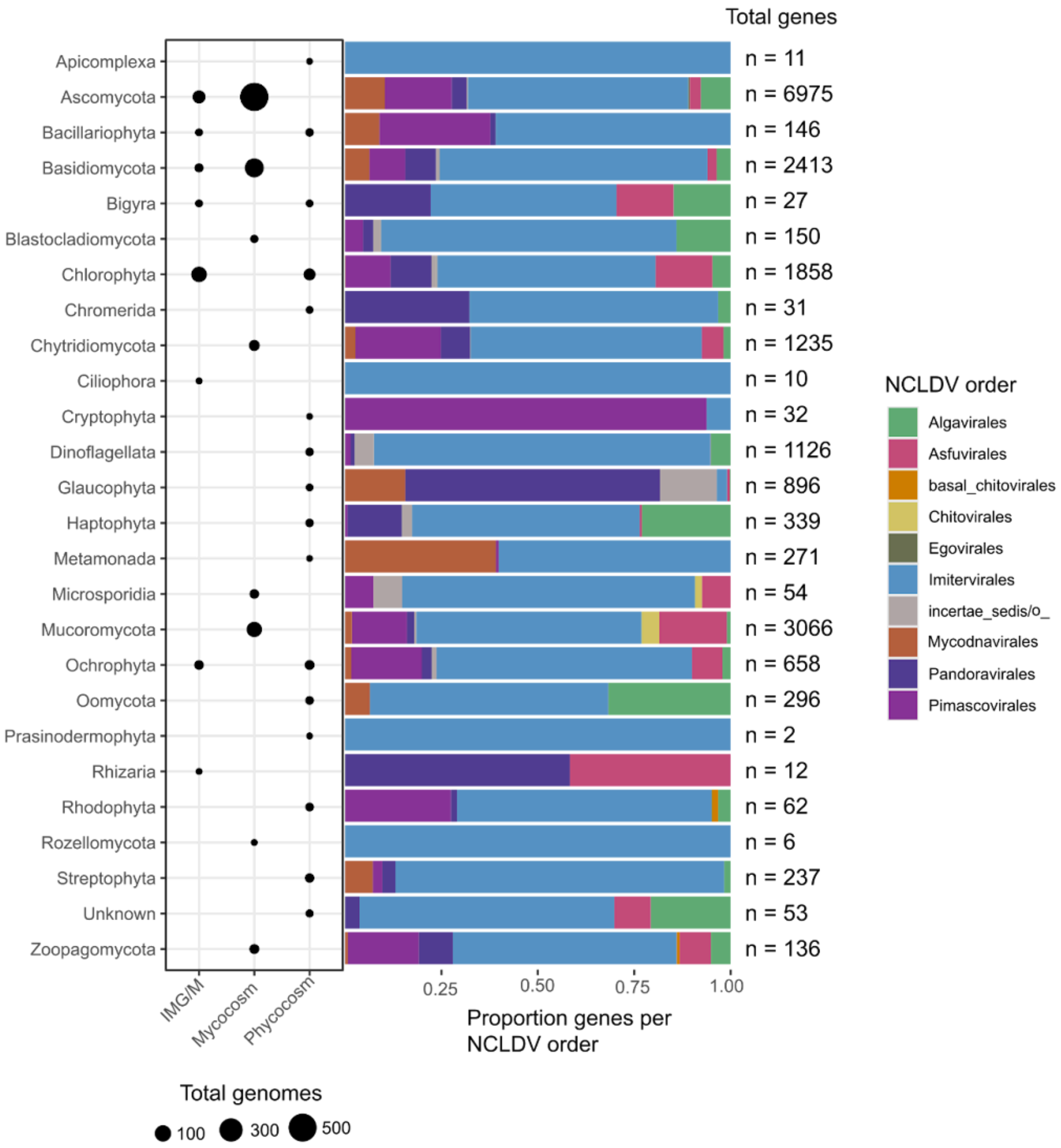


Supplementary Figure S8: Giant endogenous viral elements (GEVEs) in Joint Genome Institute databases. The number of eukaryotic genomes with GEVEs identified using a nearest neighbor search per database is shown in a dotplot. The *Nucleocytoviricota* (NCLDV) order assignment of GEVEs and the total GEVEs per eukaryotic group identified are shown in the barplot.

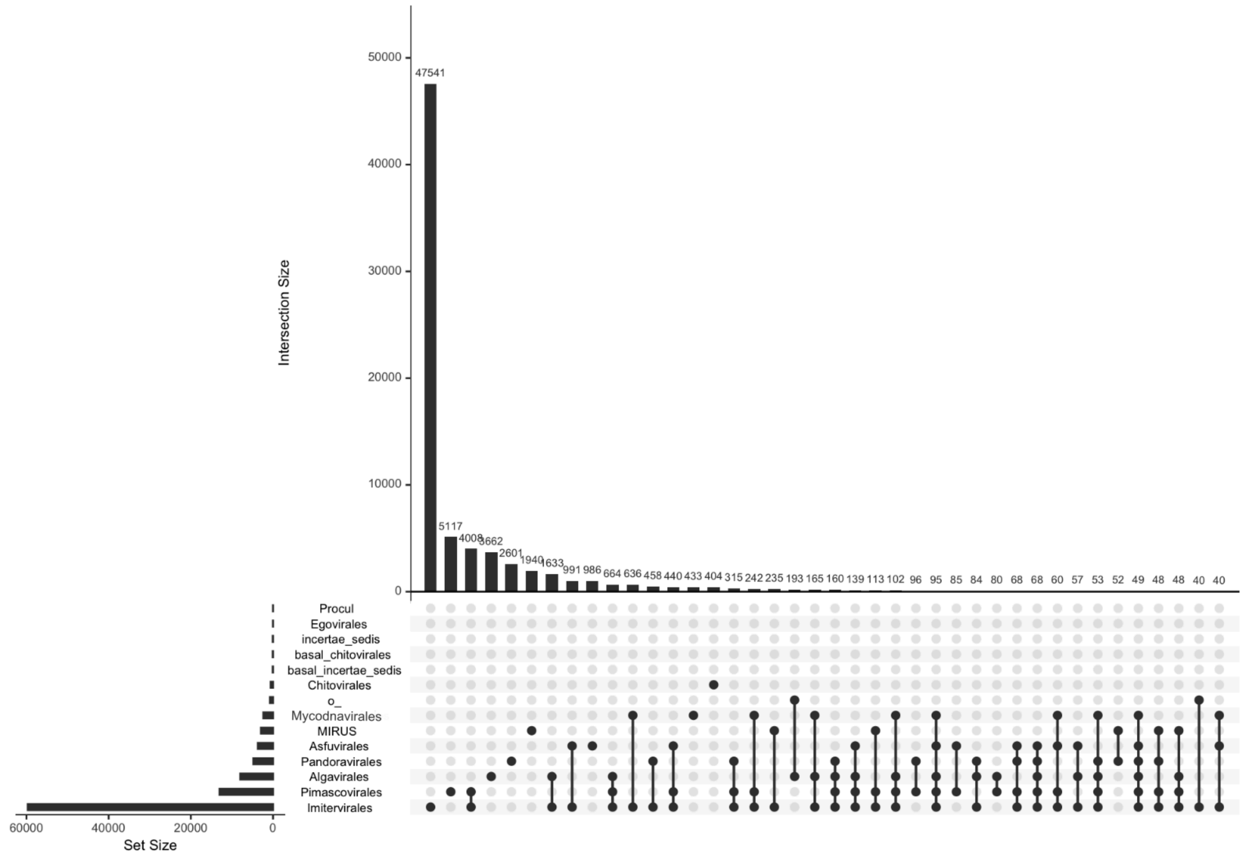


Supplementary Figure S9: Upset plot of orthogroups across orders. This plot shows the distribution and overlap of orthogroups (clusters of orthologous genes) across different orders, with the top bar chart showing intersection sizes.

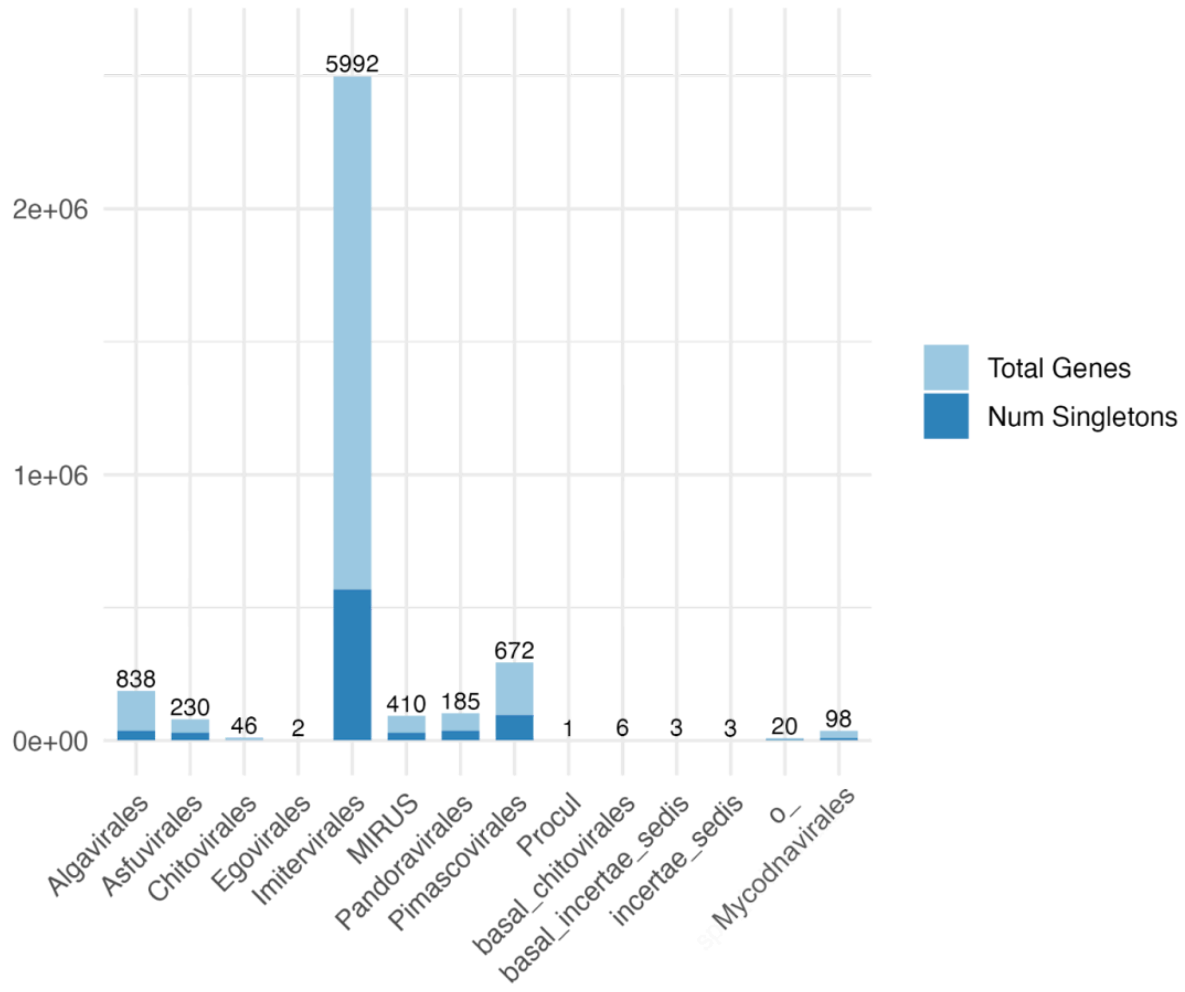


Supplementary Figure S10: Distribution of total genes and singleton genes across orders. This bar chart displays the total number of genes (light blue) and singleton genes (dark blue) for each order. Numbers on top of bar indicate the number of genomes per order.

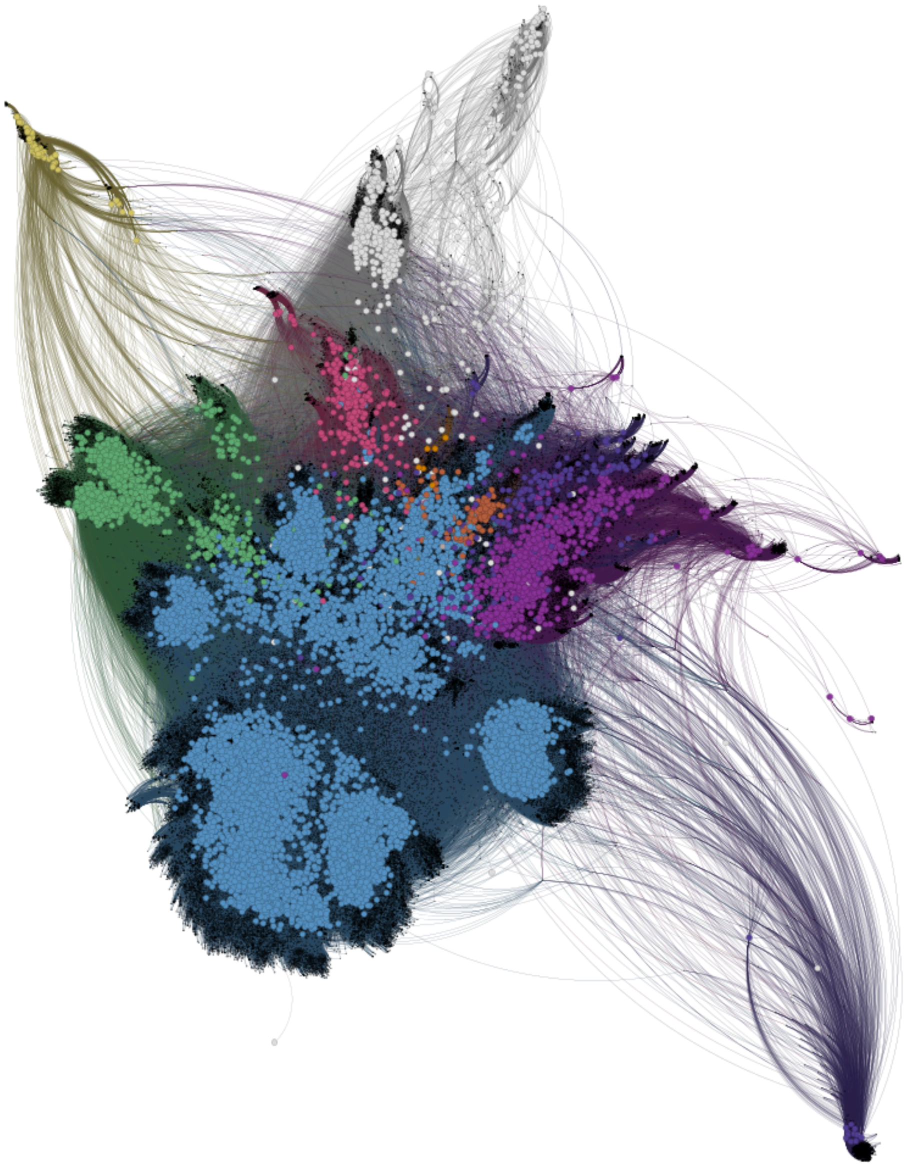

Supplementary Figure S11: Network analysis of orthogroup relationships across orders. This network visualization displays the relationships between orders based on shared orthogroups, with each colored cluster representing a different taxonomic order and edges connecting orders that share orthologous gene groups. Mirusviricota is colored by light grey.

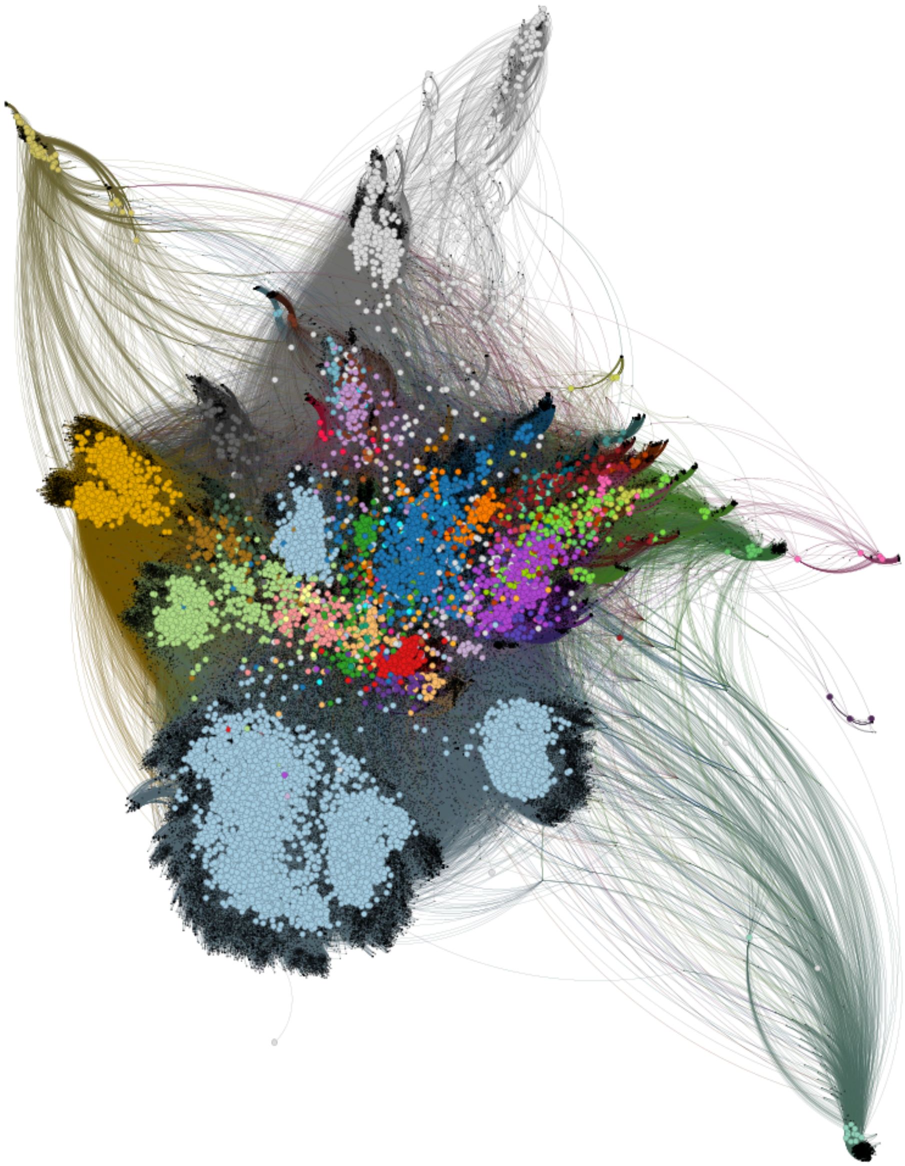

Supplementary Figure S12: Network analysis of orthogroup relationships across families. This network visualization displays the relationships between families based on shared orthogroups, with each colored cluster representing a different taxonomic family and edges connecting families that share orthologous gene groups. Mirusviricota is colored by light grey.

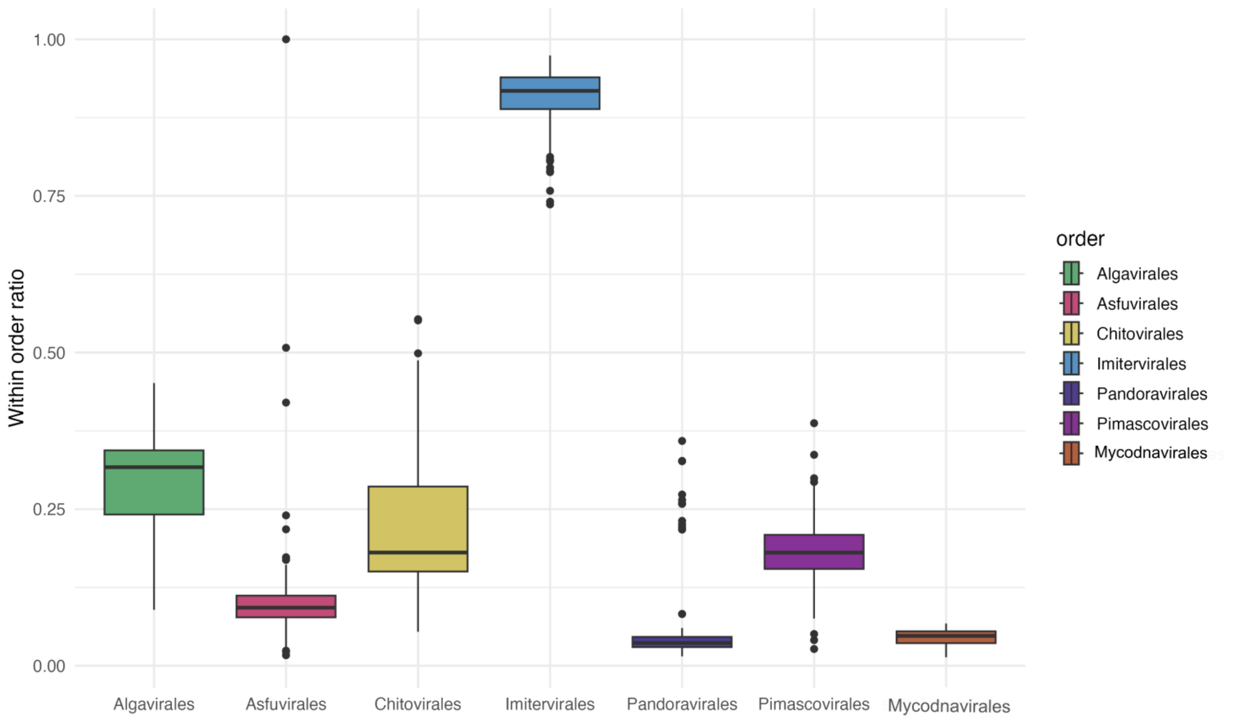


Supplementary Figure S13: Within order ratio for orthogroup network plot. This box plot displays the distribution of within-order orthogroup ratios for different orders, representing the proportion of orthogroups that are shared among members of the same order. Higher median ratio and low variability indicate extensive gene sharing within this order.

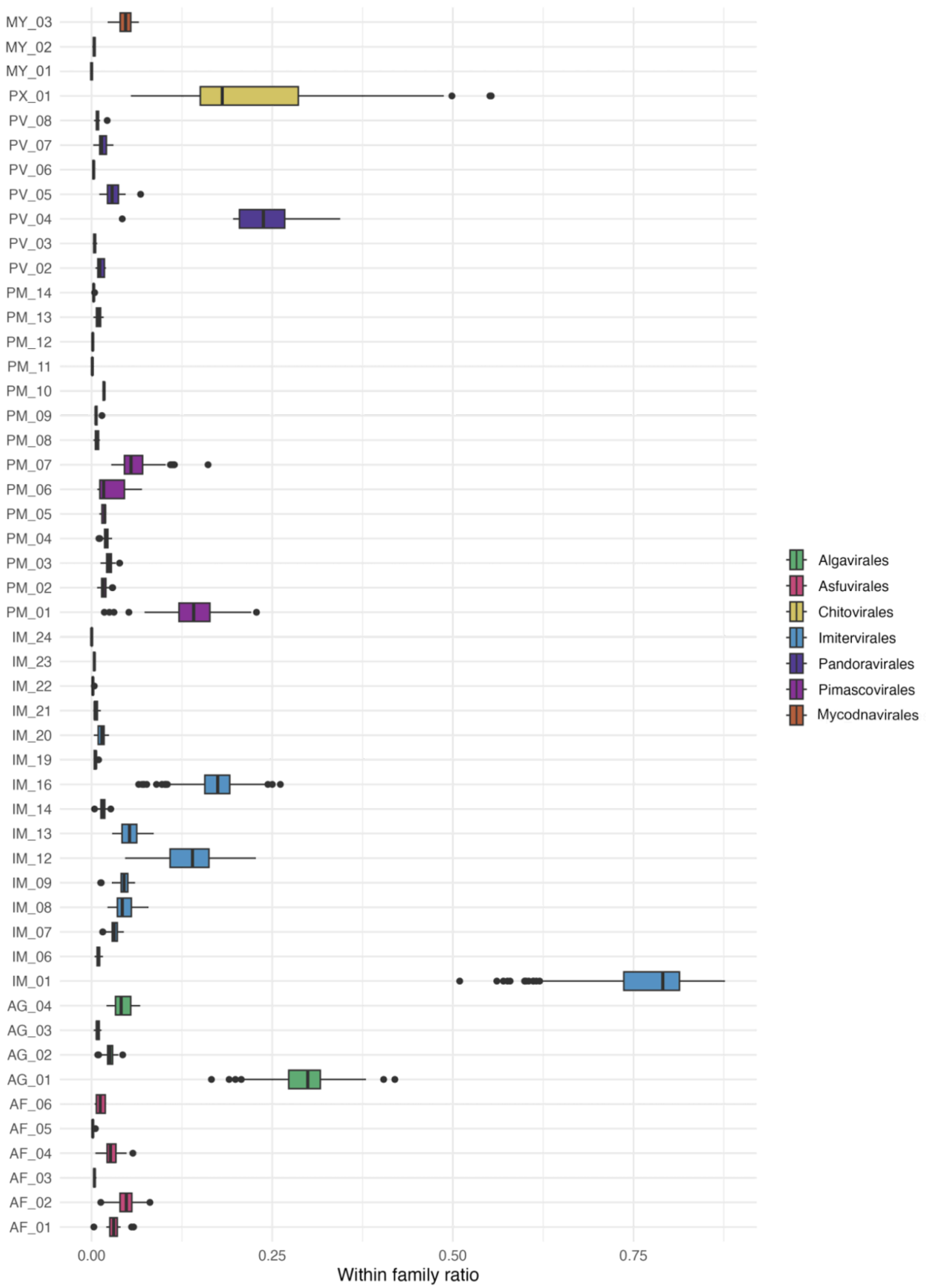

MY_03
MY_02
MY_01
PX_01
PV_08
PV_07
PV_06
PV_05
PV_04
PV_03
PV_02
PM_14
PM_13
PM_12
PM_11
PM_10
PM_09
PM_08
PM_07
PM_06
PM_05
PM_04
PM_03
PM_02
PM_01
IM_24
IM_23
IM_22
IM_21
IM_20
IM_19
IM_16
IM_14
IM_13
IM_12
IM_09
IM_08
IM_07
IM_06
IM_01
AG_04
AG_03
AG_02
AG_01
AF_06
AF_05
AF_04
AF_03
AF_02
AF_01
0.00
0.25
0.50
0.75
Within family ratio
Algavirales
Asfuvirales
Chitovirales
Imitervirales
Pandoravirales
Pimascovirales
Mycodnavirales

Supplementary Figure S14: Within family ratio for orthogroup network plot. This box plot displays the distribution of within-family orthogroup ratios for different families, representing the proportion of orthogroups that are shared among members of the same family. Higher median ratio and low variability indicate extensive gene sharing within this order.

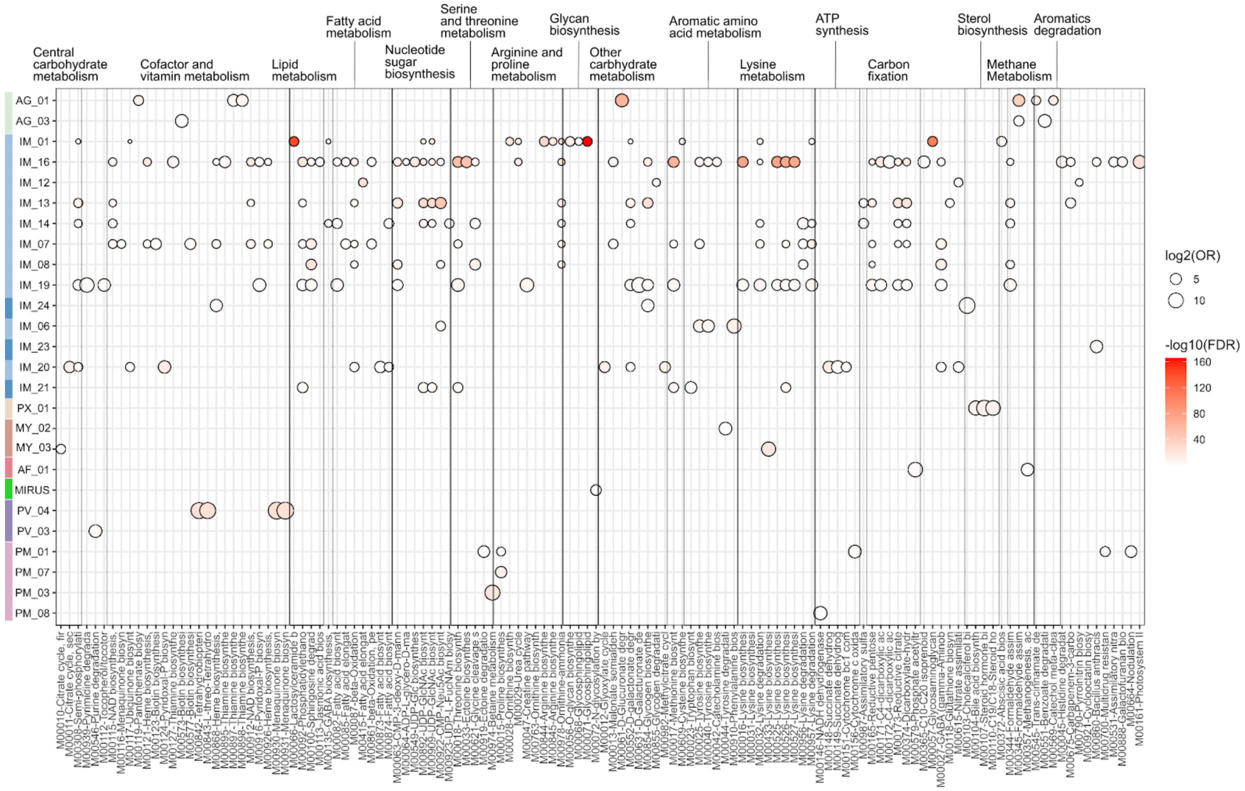


Supplementary Figure S15: Significant enrichment in modules found in only one order. An extended version, with all enriched modules, can be found at: https://yvasquez-lbl.github.io/GVMAGsV2/

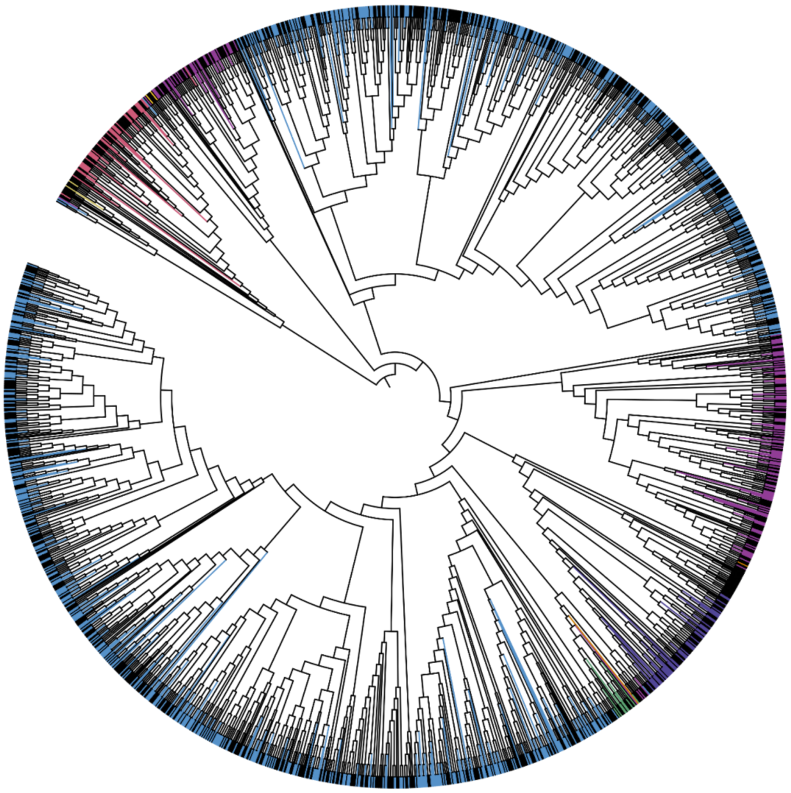

Supplementary Figure S16: Phylogenetic trees using single GVOG0022 markers with paralogs. The tree displays branches color-coded by taxonomic order.
